# Hepatic iNKT cells facilitate colorectal cancer metastasis by inducing a fibrotic niche in the liver

**DOI:** 10.1101/2024.08.19.608250

**Authors:** Marc Nater, Michael Brügger, Virginia Cecconi, Paulo Pereira, Geo Forni, Hakan Köksal, Despoina Dimakou, Michael Herbst, Anna Laura Calvanese, Giulia Lucchiari, Christoph Schneider, Tomas Valenta, Maries van den Broek

**Author notes:** Corresponding author. Winterthurerstrasse 190, 8057 Zurich, Switzerland.

## Abstract

The liver is an important metastatic organ that contains many innate immune cells, yet little is known about their role in anti-metastatic defense. We investigated how invariant natural killer T (iNKT) cells influence colorectal cancer-derived liver metastasis using different models in immunocompetent mice. We found that hepatic iNKT cells promote metastasis by creating a supportive niche for disseminated cancer cells. Mechanistically, iNKT cells respond to disseminating cancer cells by producing the fibrogenic cytokines IL-4 and IL-13 in a TCR-independent manner. Selective abrogation of IL-4 and IL-13 sensing in hepatic stellate cells prevented their transdifferentiation into extracellular matrix-producing myofibroblasts, which hindered metastatic outgrowth of disseminated cancer cells. This study highlights a novel tumor-promoting axis driven by iNKT cells in the initial stages of metastasis.

## INTRODUCTION

Colorectal cancer (CRC) is the third most common cancer worldwide. The liver is a frequent site of metastasis due to its direct connection with the gut, its low blood flow and unique vascular structure (*1–3*). In fact, liver metastases are found in 30-50% of CRC patients, often already at the time of diagnosis (*4, 5*).

A study of nearly 1000 CRC patients identified liver fibrosis as a prognostic parameter for liver metastases (*6, 7*). Fibrosis is a result of protracted inflammation and is characterized by excessive extracellular matrix (ECM) deposition (*8*). Functionally, fibrosis is thought to support malignant progression by fostering a growth-permissive environment for disseminated cancer cells (*9*). Hepatic stellate cells (HSCs) are central mediators of the fibrotic response. Upon activation, HSCs transdifferentiate into fibrogenic myofibroblasts that drive ECM accumulation and tissue remodeling (*10, 11*). HSCs can be activated by a plethora of stimuli, including oxidative stress, metabolic alterations, and inflammatory cytokines such as TGF-β, IL-4 and IL-13 (*12*).

The liver is an important immune organ that constantly faces toxins from intestinal bacteria and circulating antigens. Therefore, various mechanisms that suppress adaptive immunity are in place to maintain homeostasis (*13*). The liver contains an abundant population of resident innate immune cells, including natural killer (NK) cells, innate lymphocytes (ILC) and natural killer T (NKT) cells. The latter recognize glycolipid antigens presented by the major histocompatibility complex (MHC) class I-like CD1d molecule (*14*). CD1d-restricted NKT cells comprise two main subsets: Type I and type II NKT cells. In mice, type I NKT cells express the invariant T cell receptor (TCR) Va14-Ja18 and are therefore referred to as invariant NKT (iNKT) cells. Experimentally, iNKT cells are usually activated by α-galactosylceramide (α-GalCer), a product of marine sponges (*15*), whereas the physiological lipid antigens may be manifold and are less characterized (*14*). Type II NKT cells express a polyclonal TCR repertoire and recognize self and microbial lipids (*16*). In the mouse liver, 30% of all lymphocytes are NKT cells, most of which are iNKT cells (*17*). Mucosal-associated invariant T (MAIT) cells are thought to be the iNKT counterpart in the human liver (*18*). NKT cells patrol liver sinusoids and respond to various stimuli including TCR engagement, cytokines, and ligation of activating receptors such as Toll-like receptors (TLRs) or NKG2D (*19*, *20*). Because of their ability to produce pro- and anti-inflammatory cytokines, studies describing the influence of NKT cells on various liver pathologies reported conflicting conclusions (*21*, *22*). That NKT cells promote liver fibrosis, however, became unequivocally clear from different models of acute and chronic liver damage (*23–26*). Although fibrosis is associated with metastasis in human cancer, and the fibrogenic capacity of iNKT cells is recognized, a direct link between these processes has not yet been shown.

Here, we sought to investigate whether and how hepatic iNKT cells influence the development of liver metastasis. Using iNKT cell-deficient mouse strains and two different models for CRC-derived liver metastasis, we discovered that iNKT cells promote the outgrowth of disseminated cancer cells in the liver. Mechanistically, disseminated cancer cells activate iNKT cells to produce IL-4 and IL-13, which are sensed by HSCs and results in their activation. This results in the formation of a fibrotic niche in the liver allowing the outgrowth of disseminated cancer cells.

## RESULTS

### iNKT cells promote colorectal cancer-derived liver metastasis

To study the influence of iNKT cells on metastatic liver disease, we established two different models based on colorectal cancer (CRC) organoids. The organoids (termed APTKA (*27–29*)) harbor deletions of the genes *Apc*, *Tp53* and *Tgfbr2*, the activating mutation *Kras^G12D^*, and a myristoylated version of human *AKT.* The first model (**fig. S1A**) involves the orthotopic injection of CRC organoids into syngeneic C57BL/6 mice using an endoscopy-guided injection method (*30*, *31*). Cells from the primary colon tumor spontaneously disseminate to the liver and form macrometastases within 5-6 weeks. In the second model (**fig. S1B**), APTKA organoids are injected into the spleen and enter the portal vein, resulting in rapid seeding to the liver. The advantage of the orthotopic model is that all steps of the metastatic cascade can be studied, the disadvantage is insufficient synchronization of metastatic seeding. In contrast, the intrasplenic model enables studying the early effects of metastatic seeding and rules out a potential influence of the primary tumor on processes in the liver. Thus, we use each of the complementary models to address specific questions.

We orthotopically injected APTKA organoids into *Cd1d* ^-/-^ *and Traj18* ^-/-^ mice and used C57BL/6 mice as control. C*d1d* ^-/-^ mice lack both type I and II NKT cells, whereas *Traj18* ^-/-^ mice only lack iNKT (type I NKT) cells but still contain a sparse population of type II NKT cells (**fig. S1, C and D**) (*32*, *33*). We collected and analyzed primary colon tumors and livers 42 days after injection (**Fig. 1A**). While the weight of the primary colon tumor was similar in all strains, the metastatic rate and burden were significantly lower in iNKT cell-deficient mice (**Fig. 1, B and C**). We saw a similar protection against liver metastasis in *Rag1* ^-/-^ mice, which are devoid of NKT, T and B cells (**fig. S1, E to G**), suggesting that NKT cells do not promote metastasis by inhibiting protective effects of B or T cells.

**Fig. 1.**
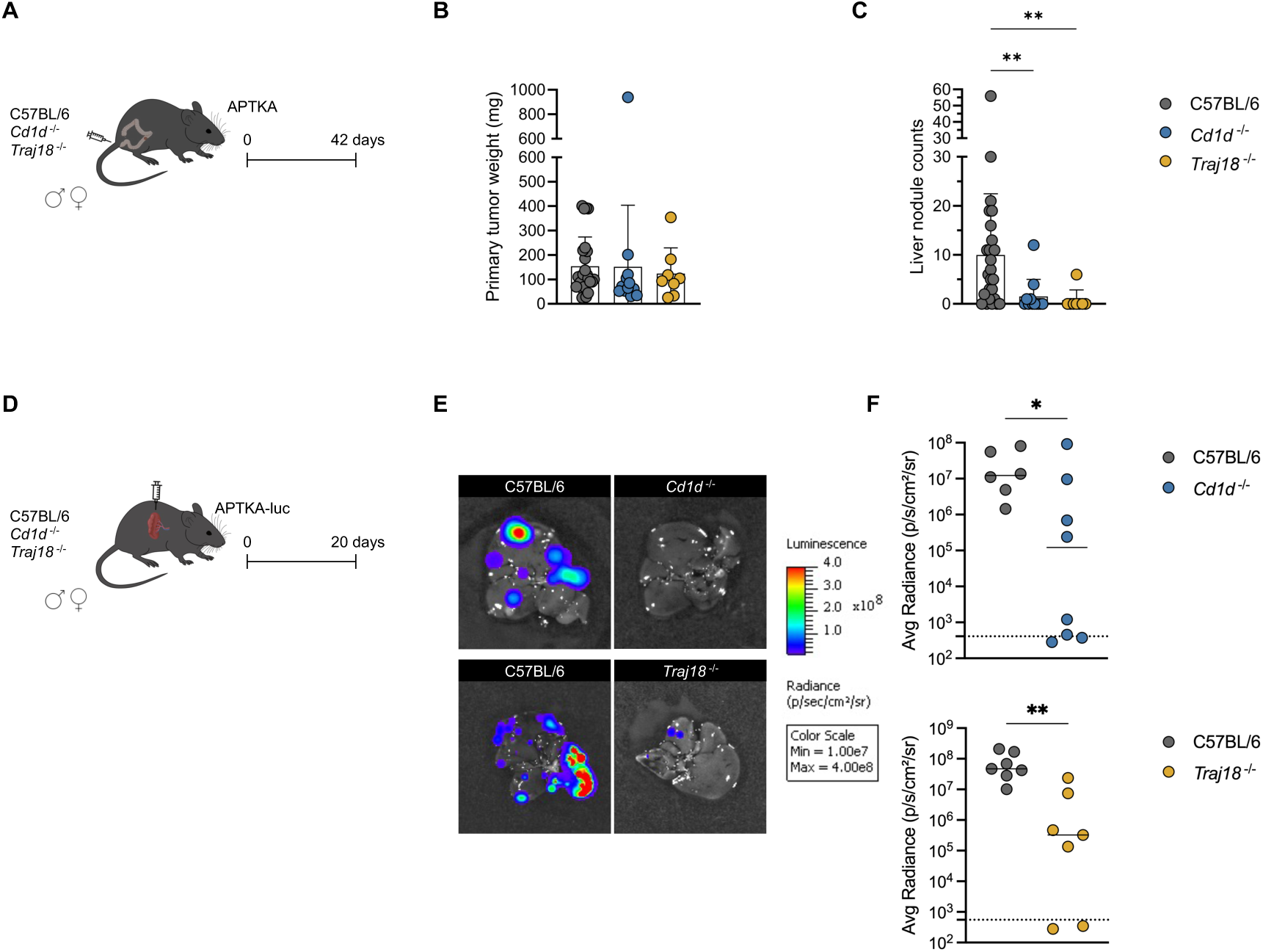
iNKT cells promote colorectal cancer-derived liver metastasis. (A) APTKA organoids were orthotopically injected into male and female C57BL/6, *Cd1d* ^-/-^ and *Traj18 ^-^*^/-^ mice. Primary tumors and livers were collected and analyzed 42 days later. (B) Weight of primary colon tumors. (C) Quantification of liver metastasis. Pooled data from 3 experiments are shown in panels B and C. Bars represent means ± SD. Symbols represent values from individual mice. \*\**p* < 0.01 (one-way ANOVA) (D) APTKA-luc organoids were intrasplenically injected into male and female C57BL/6, *Cd1d* ^-/-^ and *Traj18 ^-^*^/-^ mice. *Ex vivo* bioluminescence of the liver was measured 20 days later. (E) Representative measurement of liver bioluminescence and (F) quantification of liver bioluminescence. The dotted lines represent the limit of detection. The solid lines represent the median. Symbols represent values from individual mice. \**p* < 0.05, \*\**p* < 0.01 (unpaired Mann-Whitney U test).

To exclude systemic effects of the primary tumor, we intrasplenically injected APTKA-luc cells into *Cd1d* ^-/-^, *Traj18* ^-/-^, and C57BL/6 mice and measured the metastatic load *ex vivo* by bioluminescence 20 days later (**Fig. 1D**). This experimental setup confirmed the results of the orthotopic model (**Fig. 1, E and F**), indicating that hepatic iNKT cells promote liver metastasis *in situ* and independently of the primary tumor. To exclude that our findings are a peculiarity of APTKA organoids, we intrasplenically injected luciferase-expressing MC38 colorectal (MC38-luc) cancer cells (**fig. S1H**) and obtained similar results (**fig. S1, I and J**). Together, these results indicate that iNKT cells promote liver metastasis derived from colon cancer.

### Cancer-cell-induced hepatic fibrosis depends on iNKT cells

Fibrosis is characterized by ECM deposition and remodeling (*8*). In the liver, fibrosis is associated with progression of liver cancer (*34*), the formation of the metastatic niche (*35*, *36*) and relies on the activation of hepatic stellate cells into collagen-producing, α-SMA^+^ myofibroblasts (*11*). Using different models for liver damage, unconventional T cells were shown to be important mediators of fibrogenesis (*23–26*, *37–39*). Histological analysis of livers in the orthotopic model (**Fig. 2A**) showed increased fibrosis in metastatic lesions reflected by prominent collagen deposition (**Fig. 2B**) and the abundance of α-SMA^+^ myofibroblasts (**Fig. 2C**). In isolated metastatic nodules, we found an increased expression of *Acta2* and *Col1a1*, two transcripts associated with the activation of hepatic stellate cells (**Fig. 2, D and E**). Using *Ncr1 ^Tdtomato^ Cxcr6 ^Gfp^* reporter mice, we saw an accumulation of NKT cells (defined as CXCR6^+^ NKp46^-^ CD8^-^) in the metastatic lesions (**Fig. 2F**).

**Fig. 2.**
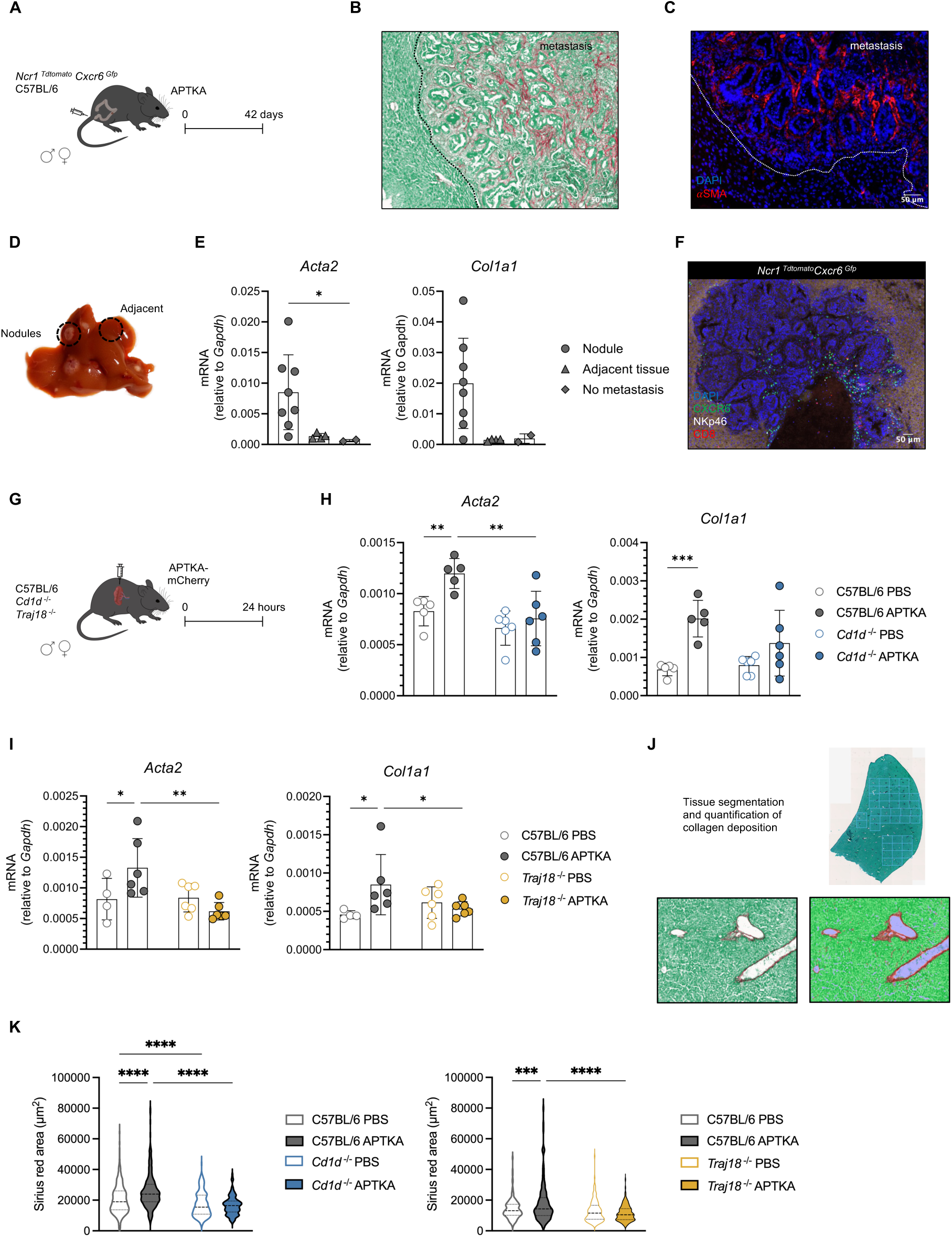
Cancer-cell-induced hepatic fibrosis depends on iNKT cells. (A) APTKA-mCherry organoids were orthotopically injected into male and female C57BL/6 or *Ncr1 ^Tdtomato^ Cxcr6 ^Gfp^* mice. Livers were collected and analyzed 42 days later. (B) Representative metastatic lesion in a C57BL/6 mouse stained with Sirius red. (C) Representative metastatic lesion in a C57BL/6 mouse stained with DAPI and anti-αSMA. (D) Parts of the metastatic livers were isolated for qRT-PCR analysis. (E) Relative quantification of *Acta2* and *Col1a1* transcripts. Bars represent means ± SD, symbols represent values from individual isolated liver parts. \**p* < 0.05 (one-way ANOVA). (F) Representative immunofluorescence image of a metastatic lesion in an *Ncr1 ^Tdtomato^ Cxcr6 ^Gfp^* mouse stained with DAPI and anti-CD8. (G) APTKA-mCherry organoids or PBS were intrasplenically injected into male and female C57BL/6, *Cd1d* ^-/-^ and *Traj18 ^-^*^/-^. Livers were collected and analyzed 24 hours later. (H) Relative quantification of *Acta2* and *Col1a1* transcripts in C57BL/6 and *Cd1d* ^-/-^ mice. (I) Relative quantification of *Acta2* and *Col1a1* transcripts in C57BL/6 and *Traj18 ^-^*^/-^ mice. Bars represent means ± SD, symbols represent values from individual mice. \**p* < 0.05, \*\**p* < 0.01 (two-way ANOVA). (J) Sirius red-stained section of liver and tissue segmentation of Sirius red-positive area. (K) Quantification of Sirius red-positive area. \*\*\**p* < 0.001, \*\*\*\**p* < 0.0001 (two-way ANOVA).

To test the hypothesis that iNKT cells are responsible for the induction of a fibrotic niche, we assessed HSC activation at the time of cancer cell dissemination to the liver. Thus, we injected C57BL/6 and iNKT-deficient *Cd1d* ^-/-^ and *Traj18* ^-/-^ mice with APTKA-mCherry organoids intrasplenically to synchronize cancer cell seeding of the liver (**Fig. 2G**). Twenty-four hours later, we saw the upregulation of *Acta2* and *Col1a1* transcripts and the deposition of collagen only in the presence of iNKT cells (**Fig. 2, H to K**). This indicates that HSC activation by disseminated cancer cells depends on the presence of iNKT cells. The absence of iNKT cells may change the structure of the liver or influence the entry of cancer cells into the liver parenchyma, both of which could be confounding factors precluding the interpretation of the abovementioned results. We measured the seeding 3 hours after intrasplenic injection of APTKA-luc cells and confirmed that the absence of iNKT cells did not affect the early seeding of cancer cells to the liver (**fig. S2, A to C**). Also, APTKA-mCherry cells were detected by immunofluorescence and measured at similar amounts by qRT-PCR in C57BL/6 and *Cd1d* ^-/-^ mice 24 hours after intrasplenic injection (**fig. S2, D and E**). Further, we excluded gross structural differences between C57BL/6 and *Cd1d* ^-/-^ mice by H&E staining of liver tissue from naive mice (**fig. S2F**).

Taken together, our results suggest that the arrival of disseminating cancer cells in the liver activates HSCs in an iNKT-dependent manner.

### Cancer cell dissemination induces activation of hepatic iNKT cells

iNKT cells can be activated via their T cell receptor involving antigens presented by the non-polymorphic MHC class I-like molecule CD1d. Alternatively and like NK cells, iNKT cells can be activated by innate signals including IL-12 and ligands for Toll-like receptors (TLRs) or NKG2D (*20*). To discriminate between TCR-dependent and - independent activation, we orthotopically injected APTKA organoids in C57BL/6 mice together with a CD1d-blocking antibody or PBS (**fig. S3A**). CD1d-blockade did not influence the weight of the primary colon tumor or liver metastasis, suggesting that the metastasis-promoting activity of iNKT cells does not require cognate TCR interaction (**fig. S3, B and C**). Next, we looked for signals that can stimulate iNKT cells independently of their TCR and found a higher expression of the NKG2D ligands *H60* and *Mult1* (*40*) in livers from APTKA-injected than from PBS-injected mice (**fig. S3D**). Along the same lines, NKG2D ligand-mediated activation of unconventional T cells has been recently described as a mechanism contributing to metabolic dysfunction-associated steatohepatitis (MASH) and fibrosis (*38*, *41*, *42*).

To better understand how disseminated cancer cells activate hepatic iNKT cells, we performed single-cell RNA sequencing of NKT cells sorted from livers 24 hours after intrasplenic injection of PBS or APTKA organoids (**Fig. 3A**). At steady state, most hepatic NKT cells express *Tbx21* suggesting a Th1-like polarization. This is in line with their potential to produce IL-4 and IFN-γ after TCR engagement (*17*). A small proportion of NKT cells expressed Th17 signature genes such as *Rorc*, *Blk*, *Tmem176a*, *Il22* and *Ccr6* (**fig. S3, E and F**). To evaluate differences in the global population of NKT cells between naïve and early metastatic livers, we performed gene set enrichment (GSE) analysis using the UCell algorithm for single-cell signature scoring (*43, 44*). NKT cells isolated from metastatic livers had a higher enrichment score for gene modules associated with NK cell activation (**fig. S3G**), underscoring our findings described above.

**Fig. 3.**
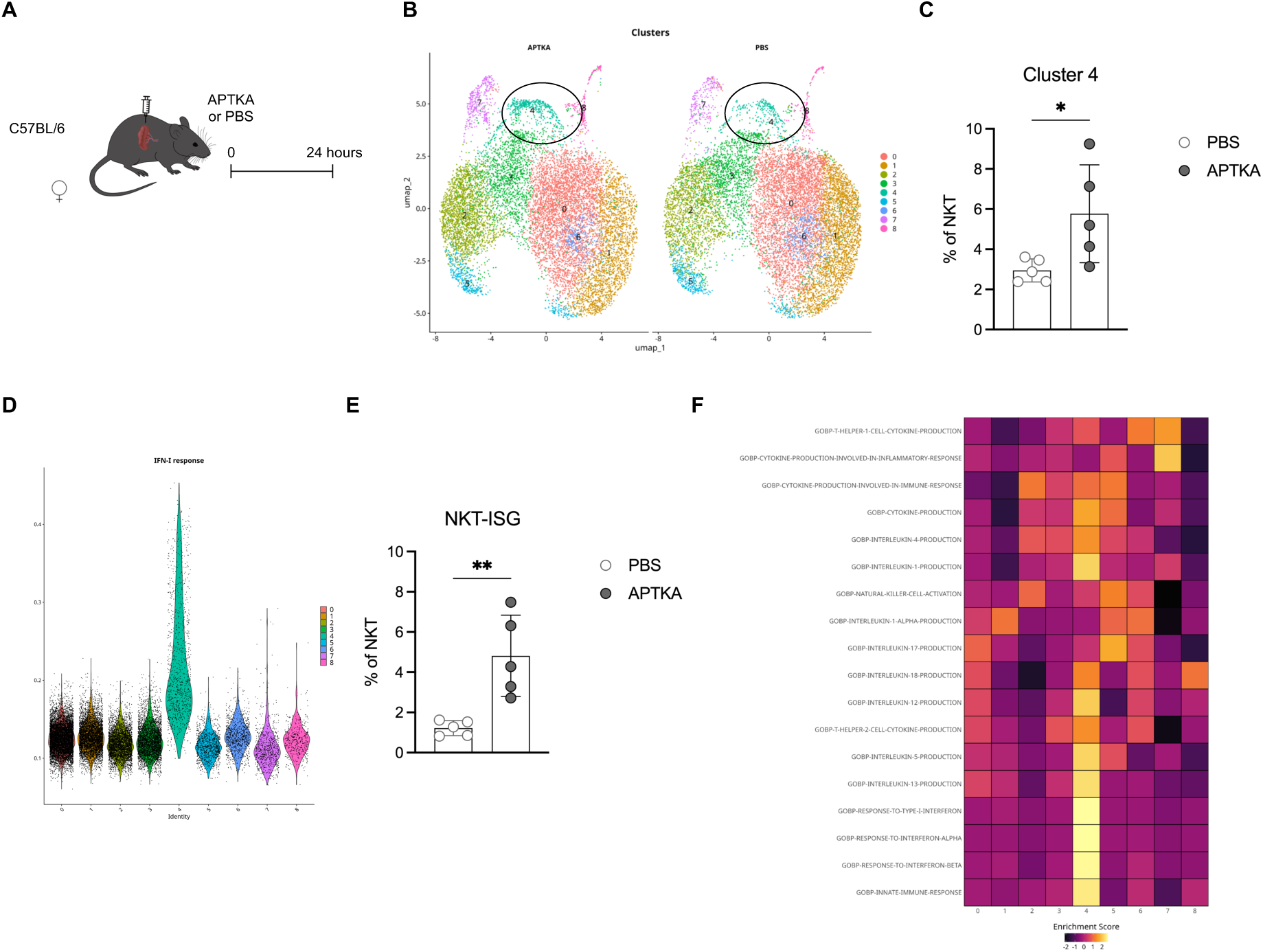
Cancer cell dissemination induces activation of hepatic iNKT cells. (A) APTKA organoids or PBS were intrasplenically injected into female C57BL/6 mice. Liver NKT cells (single, live, CD45^+^, lin^-^, CXCR6^+^, CD8^-^, NK1.1^+^, NKp46^-^ cells) were sorted 24 hours later and processed following the BD Rhapsody Single-Cell Analysis System Workflow. Sequencing was performed using the NovaSeq X Plus platform. Lin = Ly6G, CD19, CD115 and F4/80. (B) Uniform Manifold Approximation and Projection (UMAP) projection with Seurat clusters overlay. (C) Comparison of cluster 4 frequency between PBS and APTKA-injected mice. (D) Single-cell enrichment score of IFN-I response signature from UCell across identified clusters. (E) Percentage of cells with an enrichment value higher than 0.19. (F) z-transformed enrichment score of selected gene sets across identified iNKT cell clusters. Bars represent means ± SD, symbols represent values from individual mice. \**p* < 0.05, \*\**p* < 0.01 (unpaired Mann-Whitney U test).

Graph-based clustering analysis using the Louvain algorithm (*45*) identified nine distinct NKT cell clusters in the liver of which cluster 4 was uniquely enriched in the early metastatic environment (**Fig. 3, B and C**). Cluster 4 was characterized by gene signatures associated with the production of type 2 cytokines including IL-4, IL-5 and IL-13, and with IFN-I response (**Fig. 3, D to F**). The latter fits with the observed elevated abundance of *Ifnb1* in the liver lysate of APTKA-injected mice (**fig. S3H**). Together, we showed that hepatic iNKT cells are rapidly activated by disseminated cancer cells to secrete type 2 cytokines independently of TCR signaling.

### Hepatic iNKT cells respond to disseminated cancer cells by producing IL-4 and IL-13

iNKT cells can produce various cytokines upon activation, including IFN-γ, IL-4, IL-13 and TGF-β (*39*). While IFN-γ has antifibrotic potential by inducing HSC apoptosis and cell cycle arrest (*46–48*), IL-4, IL-13 and TGF-β activate HSCs *in vitro* and *in vivo* (*12*). We evaluated the production of these four cytokines in NKT cells following the early dissemination of cancer cells to the liver. We isolated RNA from NKT cells, which were sorted from the liver 24 hours after the intrasplenic injection of APTKA organoids or PBS (**Fig. 4A and fig. S4A**). Compared to NKT cells isolated from control livers, NKT cells from early metastatic livers expressed more *Il4* and *Il13* but not *Ifng* or *Tgfb1* transcripts (**Fig. 4B**). Using *Il4* ^4Get^ reporter mice, we observed an increased number of IL-4^+^ liver iNKT cells after cancer cell dissemination, thus confirming the transcript data (**Fig. 4, C to E and fig. S4B**).

**Fig. 4.**
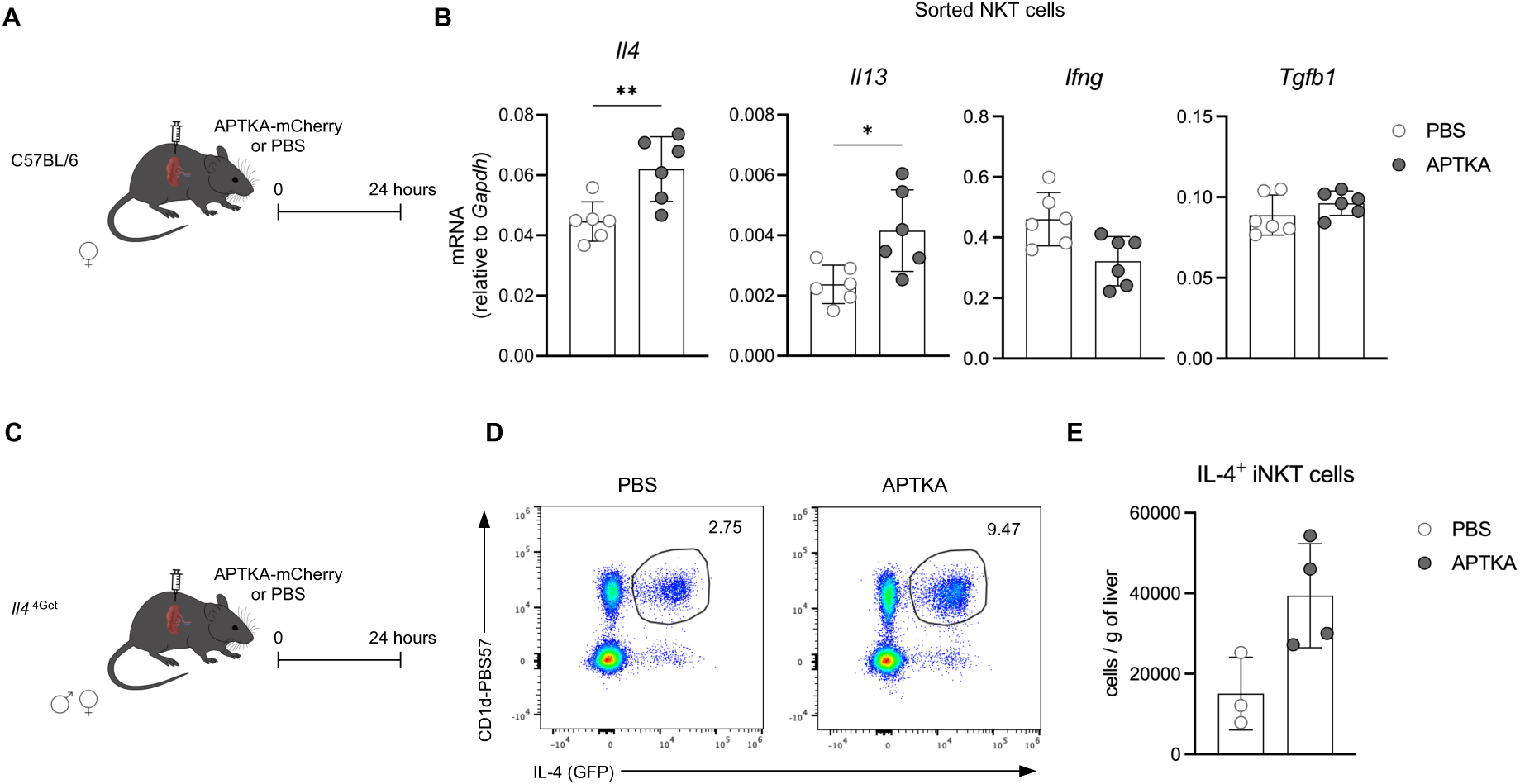
Hepatic iNKT cells respond to disseminated cancer cells by producing IL-4 and IL-13. (A) APTKA-mCherry organoids or PBS were intrasplenically injected into male and female C57BL/6 mice. NKT cells (single, live, CD45^+^, lin^-^, CXCR6^+^, CD8^-^, NK1.1^+^, NKp46^-^ cells) were sorted from livers 24 hours later and were processed for mRNA isolation. Lin = Ly6G, CD19, CD115 and F4/80. (B) Relative quantification of *Il4*, *Il13*, *Ifng*, and *Tgfb1* transcripts. (C) APTKA-mCherry organoids or PBS were intrasplenically injected into male and female *Il4* ^4Get/wt^ mice. Livers were collected 24 hours later and processed for flow cytometry. (D) Representative dot plots and (E) quantification of IL-4^+^ iNKT cells in the liver of PBS- and APTKA-injected mice. Samples were pre-gated on single, live, CD45^+^, Ly6G^-^, CD19^-^, TCRβ^+^, CD11b^-^ cells. Bars represent means ± SD, symbols represent values from individual mice. \**p* < 0.05, \*\**p* < 0.01 (unpaired Mann-Whitney U test).

Together, these results indicate that hepatic iNKT cells respond to the arrival of disseminating cancer cells by increased production of the fibrogenic cytokines IL-4 and IL-13.

### Disruption of IL-4R**α** signaling in hepatic stellate cells impairs fibrogenesis and diminishes metastatic growth in the liver

To investigate whether IL-4 and IL-13 directly activate HSC during the early phase of metastasis, we used *Lrat* ^Cre^ *Il4ra* ^fl^ mice that have an HSC-specific knockout of the IL-4Rα, which is the common subunit of the IL-4 and IL-13 receptor (*49, 50*). Following intrasplenic injection of APTKA-mCherry organoids or PBS, we observed impaired HSC activation and collagen deposition in mice with HSC-selective deficiency of IL-4Rα (**Fig. 5, A to D**). These data indicate that iNKT-derived IL-4 and IL-13 must be sensed by HSCs for their activation and subsequent initiation of fibrosis. Consequently, we expected that compromised HSC activation and diminished fibrosis result in decreased liver metastasis. To test this hypothesis, we orthotopically injected APTKA organoids in *Lrat* ^Cre^ *Il4ra* ^fl^ mice and control littermates. Five weeks later, we assessed the development of the primary tumor and liver metastasis (**Fig. 5E**). While the weight of the primary colon tumor was unaffected by HSC-specific deficiency of IL-4Rα, the incidence of liver metastasis was significantly reduced (**Fig. 5, F to H**). Thus, these findings indicate that iNKT cell-derived IL-4 and IL-13 must be sensed by HSC to prepare a fibrotic niche that promotes metastatic outgrowth of disseminated cancer cells in the liver (**Fig. 5I**).

**Fig. 5.**
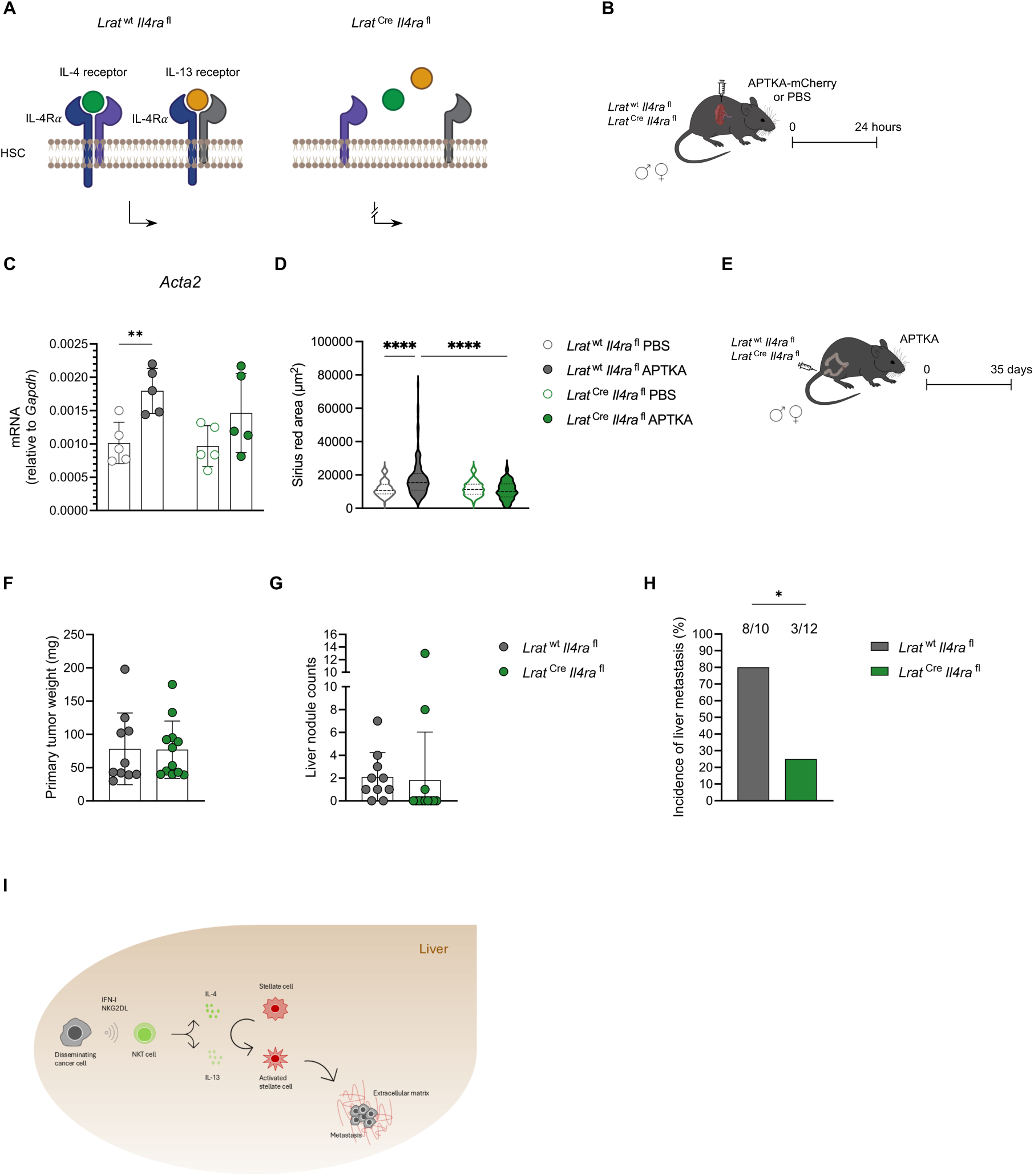
Disruption of IL-4Rα signaling in hepatic stellate cells impairs fibrogenesis and diminishes metastatic growth in the liver. (A) Scheme explaining the genetic model used. (B) APTKA-mCherry organoids were intrasplenically injected into male and female *Lrat* ^Cre/wt^ *Il4ra* ^fl/fl^ and *Lrat* ^wt/wt^ *Il4ra* ^fl/fl^ mice. Livers were collected 24 hours later. (C) Relative quantification of *Acta2* transcript. (D) Quantification of Sirius red-positive area. Bars represent means ± SD, symbols represent values from individual mice \*\**p* < 0.01, \*\*\*\**p* < 0.0001 (two-way ANOVA). (E) APTKA organoids were orthotopically injected into male and female *Lrat* ^Cre/wt^ *Il4ra* ^fl/fl^ and *Lrat* ^wt/wt^ *Il4ra* ^fl/fl^ mice. Primary tumors and livers were collected and analyzed 35 days later. (F) Weight of primary colon tumor. (G) Quantification of liver metastasis. (H) Incidence of liver metastasis in tumor-bearing mice. Bars represent means ± SD, symbols represent values from individual mice. \**p* < 0.05 (Fisher’s exact test). (I) Schematic summary of the main findings of the study.

## DISCUSSION

NKT cells are tissue-resident innate immune cells that patrol the liver sinusoids; as such, they are ideally positioned to act as immediate responders to disseminated cancer cells. The role of NKT cells in cancer is controversial. Some studies have shown tumor-restricting effects of α-GalCer-activated NKT cells in models of experimental metastasis (*51–54*) and in genetic models of solid tumors (*55*). Along the same lines, iNKT cell-deficient mice were more susceptible to methylcholanthrene-induced fibrosarcoma. Together, these results suggest that iNKT cells restrict cancer, even in the absence of exogenous stimulation (*56*). In contrast, recent work has reported tumor-promoting functions of NKT cells in primary liver cancer and metastasis (*57–61*). Those conflicting findings may be attributed to several factors. First, in many studies that showed a tumor-restricting function of NKT cells, NKT cells were stimulated with α-GalCer. Because α-GalCer binds with high affinity to CD1d and is a strong TCR agonist for iNKT cells, it has been used as a prototype stimulus for iNKT cells that triggers IFN-γ secretion. By using α-GalCer, the response of iNKT cells is skewed towards IFN-γ production and thus underestimates the effects of iNKT cells through the production of type 2 cytokines, hedgehog ligands or osteopontin (*25, 62*). We found that hepatic NKT cells, upon sensing disseminated cancer cells, did not produce IFN-γ but rather the fibrogenic cytokines IL-4 and IL-13. This suggests that using a strong stimulus such as α-GalCer to study the role of iNKT cells in liver metastasis may confound the results. Second, early work on iNKT cells used the first generation of the *Traj18* ^-/-^ strain, which was later shown to have an impaired T cell repertoire (*63*). This limitation was addressed in subsequent generations of the strain, which were developed to eliminate the confounding factors present in the initial version. (*64–66*). Therefore, data collected using the first generation of *Traj18* ^-/-^ mice to study the role of iNKT cells should be cautiously interpreted. Later studies, including ours, used the second generation of *Traj18* ^-/-^ strains. Third, the tissue context and environmental cues influence the function of NKT cells. For instance, it was shown that microbial exposure dictates whether NKT cells are pathogenic or protective in conditions such as ulcerative colitis or liver cancer (*67, 68*).

A recent study has shown that tissue-resident iNKT17 cells facilitate cancer cell extravasation into the liver via IL-22 (*19*). Whether this mechanism is generally operative in liver metastasis is currently unknown. Also, due to the unavailability of tools that target iNKT or NKT cells, mechanistic studies on how these cells influence liver metastasis are largely lacking.

We investigated whether and how iNKT cells influence the development of hepatic metastasis using two mechanistically different models for CRC-derived liver metastasis and two mouse strains that lacked either all NKT cells (*Cd1d ^-/-^*) or only iNKT cells (*Traj18 ^-/-^*). We found that in the absence of iNKT cells, disseminated cancer cells did not survive and progress to metastasis despite equal seeding of the liver. These findings prompted the hypothesis that arriving cancer cells are sensed by iNKT cells leading to their activation. To test this hypothesis, we characterized iNKT cells isolated from early metastatic and naïve livers by single-cell RNA sequencing. We found that disseminated cancer cells induced a signature of innate activation in iNKT cells, including signaling of type I interferon signaling and NKG2D ligands. Recent findings have indicated hepatocytes as the major source of NKG2D ligands in the inflamed liver. Sensing of NKG2D ligands by innate-like T cells exacerbated liver fibrosis in a model of MAFLD (*38*). Besides innate activation signatures, we found an increase in IL-4 and IL-13 within a day after sensing cancer cells.

These two type 2 cytokines raised our interest because of their established role in liver fibrosis (*70*). We show that IL-4 and IL-13 directly activate HSCs, leading to their transdifferentiation into myofibroblasts and subsequent deposition of extracellular matrix. HSCs were recently identified as the primary source of myofibroblasts in the liver and as key players in fibrosis (*11*). Depletion of HSCs showed their essential role in metastatic outgrowth in the liver (*71*). The underlying mechanisms, however, were not addressed in this study and remain largely unexplored. We discovered here that HSC-specific ablation of the IL4-Rα, which binds both IL-4 and IL-13, significantly impaired fibrosis and reduced the metastatic burden, underscoring the crucial role of the IL4-Rα-signaling pathway in the establishment of the metastatic niche. Our findings are in line with studies aiming to identify fibrotic drivers in type 2-driven disease. Indeed, *Il4ra* ^-/-^ mice were protected against IL-33-driven fibrosis (*72*). Further, ablating IL4-Rα signaling in PDGFRB^+^ fibroblasts, but not in albumin-expressing hepatocytes or cholangiocytes protected mice from liver fibrosis driven by IL-13 overexpression or *Schistosoma mansoni* infection (*73*).

In conclusion, by establishing the mechanistic link between iNKT cells, fibrosis and liver metastasis, we discovered an IL-4/IL-13-driven tumor-promoting function of iNKT cells in the early stages of liver metastasis.

## MATERIALS AND METHODS

### Study Design

We aimed to elucidate the role of invariant natural killer T (iNKT) cells in the development of colorectal liver metastasis. To achieve this, we used two complementary liver metastasis models based on the injection of CRC organoids either orthotopically into the colon or intrasplenically. In the orthotopic model, mice that did not develop a primary colon tumor were excluded from the analysis. Our experimental readouts included flow cytometry, histology and immunofluorescence, single-cell RNA sequencing, and quantitative real-time PCR (qRT-PCR). Sample sizes were determined by a power analysis based on pilot studies, and each experiment was performed at least twice. All experiments were conducted in accordance with the 3R principles (Replacement, Reduction, Refinement) to minimize animal use.

### Mice

C57BL/6NRj mice were purchased from Janvier Labs. *Cd1d* ^-/-^ mice (*74*) and *Cxcr6 ^Gfp/Gfp^* mice (*75*) were purchased from The Jackson Laboratory. *Traj18* ^-/-^ mice (*66*) were provided by Paolo Dellabona (IRCCS, Italy). R26R ^Ai14/Ai14^ mice (*76*) were provided by Melanie Greter (UZH, Switzerland). *Ncr1* ^Cre/wt^ R26R ^Ai14/wt^ mice (*77*) were provided by Sonia Tugues (UZH, Switzerland). *Ncr1* ^Cre/wt^ R26R ^Ai14/wt^ were crossed to *Cxcr6 ^Gfp^*^/*Gfp*^ mice to obtain *Ncr1* ^Cre/wt^ R26R ^Ai14/wt^ *Cxcr6* ^Gfp/wt^ (*Ncr1 ^Tdtomato^ Cxcr6 ^Gfp^*) mice. *Lrat* ^Cre/wt^ R26R ^Ai14/Ai14^ mice (*11*) were provided by Ingmar Mederacke (MHH, Germany). *Il4ra* ^fl/fl^ *ll13* ^Smart13/Smart13^ *Arg1 ^Yfp/Yfp^* mice (*50, 78, 79*) were crossed to *Lrat* ^Cre/wt^ R26R ^Ai14/Ai14^ mice to obtain *Lrat* ^Cre/wt^ *Il4ra* ^fl/fl^ mice. *Lrat* ^Cre/wt^ *Il4ra* ^fl/fl^ mice were used for the selective deletion of *Il4ra* in hepatic stellate cells. Cre-negative littermates were used as controls. *Rag1* ^-/-^ mice were provided by Burkhard Becher (UZH, Switzerland). *Il4* ^4Get/wt^ mice (*80*) were provided by Manfred Kopf (ETHZ, Switzerland).

All strains have a C57BL/6 background. Eight-to-twelve-week-old mice of either sex were used with equal age- and sex-distribution among experimental groups. Breeding and experiments were performed under specific pathogen-free (SPF) conditions in facilities of the Laboratory Animal Services Center (LASC) at the University of Zurich. Mice had access to food and water *ad libitum* and were kept in a 12-hour light/dark cycle. Mouse experiments were approved by the Swiss Cantonal Veterinary Office (license numbers 156/2018 and 031/2021).

### Organoids and cancer cell lines

APTKA organoids were provided by Florian Greten (Georg-Speyer-Haus, Germany). The MC38 cell line was originally provided by Mark Smyth (QIMR Berghofer Medical Research Institute, Australia). Cells were tested negative for Mycoplasma ssp. by PCR analysis. Cells were also tested negative for 18 additional mouse pathogens by PCR (IMPACT II Test, IDEXX Bioanalytics).

Organoids and cancer cells were cultured at 37°C in a humid atmosphere with 5% CO_2_. APTKA organoids were embedded in Basement Matrix Extract type 2 (BME, Bio-Techne) and cultured with organoid medium, consisting of Advanced DMEM/F-12 (Gibco), supplemented with 10 mM HEPES (Gibco), 100 U/ml Penicillin, 100 μg/ml Streptomycin (Sigma-Aldrich), 2 mM L-Glutamine (Gibco), 1.25 mM N-Acetyl-L-cysteine (Sigma-Aldrich), 1% N-2 and 2% B-27 supplements (Gibco). MC38 cells were cultured in DMEM (Gibco) supplemented with 10% fetal bovine serum (Gibco), 100 U/ml Penicillin, 100 μg/ml Streptomycin (Sigma-Aldrich) and 2 mM L-Glutamine (Gibco).

mCherry-expressing APTKA (APTKA-mCherry), luciferase-expressing MC38 (MC38-luc) and luciferase-expressing APTKA (APTKA-luc) organoids were generated by lentiviral transduction using 8 µg/ml polybrene (Sigma-Aldrich). Lentiviruses were produced by co-transfection of the respective lentiviral expression vector pLV-mCherry (Addgene #36084) and pLenti PGK V5-LUC Neo (Addgene #21471) and the viral packaging envelope plasmids pMD2g (Addgene #12259) and pCMV-dR8.91 (provided by Christian Münz, UZH, Switzerland) into 293T cells (ATCC, CRL-3216). A DNA:polyethylenimine (Polysciences) µg ratio of 1:3 was used for transfection. mCherry^+^ cells were sorted by Fluorescence-activated cell sorting (FACS). Luciferase-expressing MC38 and APTKA cells were selected with 500 µg/ml G418 (InvivoGen) for 10 days. Clonal selection was then performed for all cell and organoid lines. The reagents used for cell and organoid culture are listed in table S1.

### Collection and expansion of APTKA organoids

Cryopreserved organoids were thawed and seeded in 4 domes of BME (75 µl each) on 6-well plates and transferred to a 37°C cell culture incubator. Once the BME was polymerized, 3 ml of organoid medium were added to each well. To collect organoids for passaging or injection, the medium was aspirated. One ml of ice-cold PBS was added per seeded well. The domes from each well were mechanically disrupted using a bent 1-ml pipette tip, and organoids were transferred into a 15-ml tube containing 10 ml of ice-cold PBS. The organoids were then centrifuged for 5 minutes at 290 g, and the supernatant was aspirated. The cell pellet was resuspended with a 200-µl bent pipette tip in 0.5 ml of ice-cold PBS to dissociate the organoids. Ten ml of ice-cold PBS was then added. The cells were centrifuged for 5 minutes at 290 g and supernatant was aspirated. For further culture, dissociated APTKA organoids were resuspended in BME.

### Orthotopic injection of APTKA organoids

Orthotopic injection of organoids was performed as described (31). Following harvesting as outlined above, two domes (approximately 1.5 x 10^5^ cells) of APTKA or APTKA-mCherry organoids were resuspended in 70 µl PBS. Mice were anesthetized and transferred onto a warming pad set at 37°C. Ophthalmic ointment (Alcon) was applied to the eyes to prevent drying. The colon was cleaned with a rapid enema with 5-10 ml PBS pre-warmed to 30°C using the plastic tubing of a 20G x 48 mm intravascular catheter (BD) attached to a 20-ml syringe (B Braun). Organoids were injected into the distal colon mucosa by colonoscopy-guided mucosal injection with a custom injection needle (Hamilton).

MC38 cells were not used in the orthotopic model due to the failure to form a primary colon tumor. The equipment used for orthotopic injection of organoids is listed in table S2.

### Intrasplenic injection of APTKA organoids or MC38 cells

Intrasplenic injection of APTKA organoids or MC38 cells was performed as described (*81, 82*). 1.5 x 10^5^ cells from dissociated APTKA-luc or APTKA-mCherry organoids or 5 x 10^5^ MC38-luc cells were resuspended in 50 µl PBS. Pre-emptive analgesia (0.04 mg/kg fentanyl and 0.1 mg/kg buprenorphine) was administered by intraperitoneal injection. Mice were anesthetized and transferred onto a warming pad set at 37°C. Ophthalmic ointment (Alcon) was applied to the eyes to prevent drying. The mice were placed in a lateral position on the right side. The fur along the torso was shaved, and the skin was scrubbed with a sterile pad wetted with 70% ethanol. A single 1-cm incision was made in the left upper abdominal wall to access the peritoneal cavity, and the spleen was exposed on a sterile gaze using surgical forceps. APTKA organoids or MC38 cells were injected into the spleen using a 0.5-ml U-100 insulin syringe with a 30G x 8 mm needle. After 15 seconds, the needle was retracted, and the injection site was compressed using a sterile gauze. After 3 minutes, the splenic artery and vein were ligated near the hilum with 2 stitches each using a surgical needle holder and absorbable coated polyglactin sutures (Ethicon). Immediately thereafter, the spleen was resected. The peritoneal wall was sutured with 3 stitches and the skin was closed with wound clips (Clay Adams). Mice received buprenorphine (0.01 mg/ml) *ad libitum* for 48 hours after surgery. The equipment used for orthotopic injection of organoids is listed in table S2.

### *Ex vivo* bioluminescence

APTKA-luc- and MC38-luc-derived metastases were visualized and quantified using an IVIS 200 imaging system (PerkinElmer) 20 minutes after intraperitoneal injection (i.p.) of 150 mg/kg D-Luciferin in PBS (Promega). The reagents and equipment used for *ex vivo* bioluminescence are listed in table S1 and table S2.

### CD1d blockade

Anti-mouse CD1d (100 μg/mouse, BioXCell) was injected i.p. in 100 µl PBS immediately after organoid injection and then twice a week. Control mice were injected with 100 µl of PBS. The reagents used for CD1d blockade are listed in table S1.

### Sorting of NKT cells

Livers were digested into a single-cell suspension. Lymphocyte enrichment was performed using a density gradient centrifugation step (Percoll, Sigma-Aldrich). The gating strategy did not include staining for TCR-β or the use of CD1d tetramer loaded with PBS-57 (an α-GalCer analog) to prevent potential transcriptional changes induced by the interaction of the TCR. Instead, NKT cells were sorted as single, live, CD45^+^, Ly6G^-^, CD19^-^, CD115^-^, F4/80^-^, CXCR6^+^, CD8^-^, NK1.1^+^ NKp46^-^ cells using a FACSAria III (BD) cell sorter. The antibodies and equipment used for sorting NKT cells are listed in table S1, table S2 and table S3.

### qRT-PCR

Fifty mg of liver tissue was homogenized in GENEzol Reagent (Geneaid) using zirconium oxide beads (Labgene Scientific) and a FastPrep tissue homogenizer (MP Biomedicals). Total RNA was isolated by phenol-chloroform extraction following the manufacturer’s instructions. 2 units of DNAse I (Thermo Fisher Scientific) per 10 µg of template RNA were used to digest genomic DNA.

RNA was extracted from sorted NKT cells using a Quick-RNA Microprep Kit (Zymo Research) following the manufacturer’s instructions.

Total RNA was reverse transcribed using a high-capacity cDNA reverse transcription kit (Applied Biosystems). Real-time quantitative PCR was performed using PowerUp SYBR Green (Applied Biosystems) or TaqMan Fast Advanced Master Mix (Applied Biosystems). *Gapdh* was used as an internal reference gene, and the relative expression of target genes was calculated as 2^−ΔCt^. The kits and reagents used for RNA isolation and cDNA synthesis are listed in table S1. The primers and probes used for qRT-PCR are listed in table S4.

### Flow cytometry

Livers were harvested in RPMI supplemented with 10% FCS and digested with collagenase IV (1 mg/ml, Thermo Fisher Scientific) and DNase I (50 µg/ml, Roche) for 45 minutes at 37°C on a rotating device. Cells were washed with PBS containing 2 mM EDTA and filtered through a 70-µm strainer (Biologix). Red blood cells were lysed using a red blood cell lysis buffer (17 mM Tris pH 7.2, 144 mM NH_4_Cl) for 2 minutes and cells were washed with PBS. Following 10 minutes of incubation with anti-CD16/32 in PBS containing 2 mM EDTA, single-cell suspensions were stained for surface molecules in 50 µl antibody mix in PBS containing 2 mM EDTA for 30 minutes at 4°C. For CD1d tetramer staining, CD1d-PBS57 or CD1d-unloaded was added to the surface antibody mix and cells were stained for 15 minutes at room temperature and then 30 minutes at 4°C. For intracellular staining, cells were fixed and permeabilized for 40 minutes at 4°C using the Foxp3/Transcription Factor Staining Buffer Set (Invitrogen). Single cell suspensions were then stained for intracellular molecules in permeabilization buffer overnight at 4°C. Samples were washed once with permeabilization buffer and once with PBS 2mM EDTA and were acquired using a Cytek Aurora spectral flow cytometer (5L). The reagents used for liver digestion are listed in table S1. The antibodies and dyes used for flow cytometry are listed in table S3.

### Immunofluorescence

Livers were fixed in 4% formaldehyde (Carl Roth) for 4 hours at room temperature and subsequently dehydrated in 15%, followed by 30% sucrose (Carl Roth) in PBS for 24 hours in each solution at 4 °C. Livers were then embedded and frozen in O.C.T. (Sakura), and 10-μm cryosections were cut using a cryotome. Sections were dried for 30 minutes at 37 °C, washed with PBS, and then blocked and permeabilized with PBS containing 4% bovine serum albumin (Carl Roth) and 0.1% Triton X-100 (Sigma-Aldrich) at room temperature for 10 minutes. Slides were then washed with PBS containing 0.05% Tween 20 (Sigma-Aldrich) and stained with a primary antibody in PBS containing 1% BSA at 4 °C overnight. Next, slides were washed and stained with a secondary antibody mix in PBS 1% BSA. Finally, slides were washed and incubated with 0.5 μg/ml DAPI (Invitrogen) for 5 minutes and mounted with Prolong Diamond medium (Invitrogen). Stained slides were scanned using the automated multispectral microscopy system Vectra 3.0 (PerkinElmer). Spectral unmixing was performed using inForm software v2.6 (PerkinElmer). The antibodies and dyes used for immunofluorescence are listed in table S3.

### Sirius red staining

Livers were fixed, dehydrated, frozen and sectioned as previously described in the “Immunofluorescence” section. Frozen liver sections were dried for 30 minutes at 37°C and fixed in Kahle fixative (4% Formaldehyde, 0.02% acetic acid and 28% ethanol in distilled H_2_O) for 10 minutes at room temperature. The fixed sections were washed in distilled H_2_O and then incubated in 0.04% Fast Green (Sigma-Aldrich) for 15 minutes at room temperature. The sections were then washed in distilled H_2_O and stained with 0.1% Fast Green/0.04% Sirius Red (Sigma-Aldrich) in saturated picric acid (Sigma-Aldrich) for 30 minutes at room temperature. Sections were washed 2 times for 5 minutes in 0.5% glacial acetic acid (Sigma-Aldrich). Tissue sections were dehydrated through an ethanol series (70%, 80%, 96% and 100% ethanol, 30 seconds each) and cleared in Ultraclear (Biosystems) for 2 minutes. Slides were mounted with Eukitt mounting medium (Sigma-Aldrich) and stored at room temperature protected from light. Stained slides were scanned using the automated multispectral microscopy system Vectra 3.0 (PerkinElmer). Tissue segmentation and quantification of Sirius red-positive area was performed on twenty images (20 x magnification) per liver and analyzed using inForm software v2.6 (PerkinElmer). The reagents used for Sirius red staining are listed in table S1.

### Single-cell RNA sequencing

Single-cell suspensions were prepared from individual liver samples (n = 5 per condition). Cells were labeled using the BD Single-Cell Multiplexing Kit and pooled according to their respective conditions.

Liver NKT cells were isolated using a BD FACSAria III cell sorter as single, live, CD45^+^, CD115^−^, F4/80^−^, CD19^−^, Ly6G^−^, CXCR6^+^, CD8^−^, NK1.1^+^, NKp46^−^ cells. 40’000 cells were loaded onto a BD Rhapsody cartridge and processed for cDNA synthesis according to the BD Rhapsody Single-Cell Capture and cDNA Synthesis protocol (Doc ID: 23-22951(01)). Library amplification was then carried out following the BD Rhapsody System mRNA Whole Transcriptome Analysis (WTA) and Sample Tag Library Preparation Protocol (Doc ID: 23-24119 (02)). Sequencing was conducted on an Illumina NovaSeq X Plus platform using 150 bp paired-end reads (R1 = 51 cycles, R2 = 71 cycles), achieving an average sequencing depth of approximately 50,000 reads per cell. Read alignment, cell barcode demultiplexing, deduplication of reads, generation of expression matrices and quality report were performed using the Rhapsody Sequence Analysis Pipeline v2.0.

The expression matrices were further analyzed in R v4.2.2. Cells were filtered to exclude those with more than 10% mitochondrial content and less than 2% ribosomal protein content. Duplicate cells were removed using scDblFinder (*83*). The Seurat pipeline v5 was employed for normalization, dimensionality reduction, and clustering of the expression data (*84*). Gene set enrichment analysis was conducted using the UCell algorithm via the escape R package v1.99 (*43, 44*). The kits and reagents used for multiplexing, cDNA synthesis and library preparation are listed in table S1.

### Statistical analysis

Statistical significance was calculated with one-way analysis of variance (ANOVA) with Dunn’s multiple comparisons correction when comparing the means of three or more independent groups. Two-way ANOVA was employed to examine the interaction between two independent variables on a dependent variable. Fisher’s Least Significant Difference (LSD) post-hoc test was used for pairwise comparisons between group means. Mann-Whitney two-tailed U tests were applied to compare the medians of two independent groups. *p* < 0.05 was considered significant (\**p* < 0.05, \*\**p* < 0.01, \*\*\**p* < 0.001, and \*\*\*\**p* < 0.0001). Statistical analysis was performed using GraphPad statistical software v10.2.0 (GraphPad Software Inc.).

## Acknowledgments

The authors thank Florian Greten (Georg-Speyer-Haus, Frankfurt am Main, Germany) for sharing the APTKA organoids, and Andreas Moor and Costanza Borrelli (ETH Zurich) for initial help with organoid handling. We thank the personnel of the Laboratory Animal Services Center (LASC, University of Zurich) for expert animal care. We thank Stephan Benke (Cytometry Facility, University of Zurich) for support with flow cytometry data analysis. We thank Tatiane Gorski (Cytometry Facility, University of Zurich) and Hubert Rehrauer (Functional Genomic Center Zurich) for help with scRNA-sequencing, and Achim Weber, Christian Münz and Sònia Tugues (University of Zurich), Matteo Iannacone (Ospedale San Raffaele, Milano, Italy), and Martin Guilliams (VIB Ghent, Belgium) for helpful discussion and suggestions.

## Author contributions

MN and MvdB conceived the experiments and wrote the manuscript; MN, MB, VC, PP, DD, MH, ALC, GF, TV and GL performed the experiments. CS provided essential reagents. MN and HK analyzed the data. MvdB secured funding; All the authors reviewed the results and approved the final manuscript.

## Data and materials availability

All data needed to obtain the conclusions in this work are present in the Supplementary Materials. All mouse lines, reagents, and software used are listed in the Methods or the Supplementary Materials. The sequencing data that were generated for this study have been deposited in Gene Expression Omnibus under the accession number GSE274890.

---

**fig. S1.**
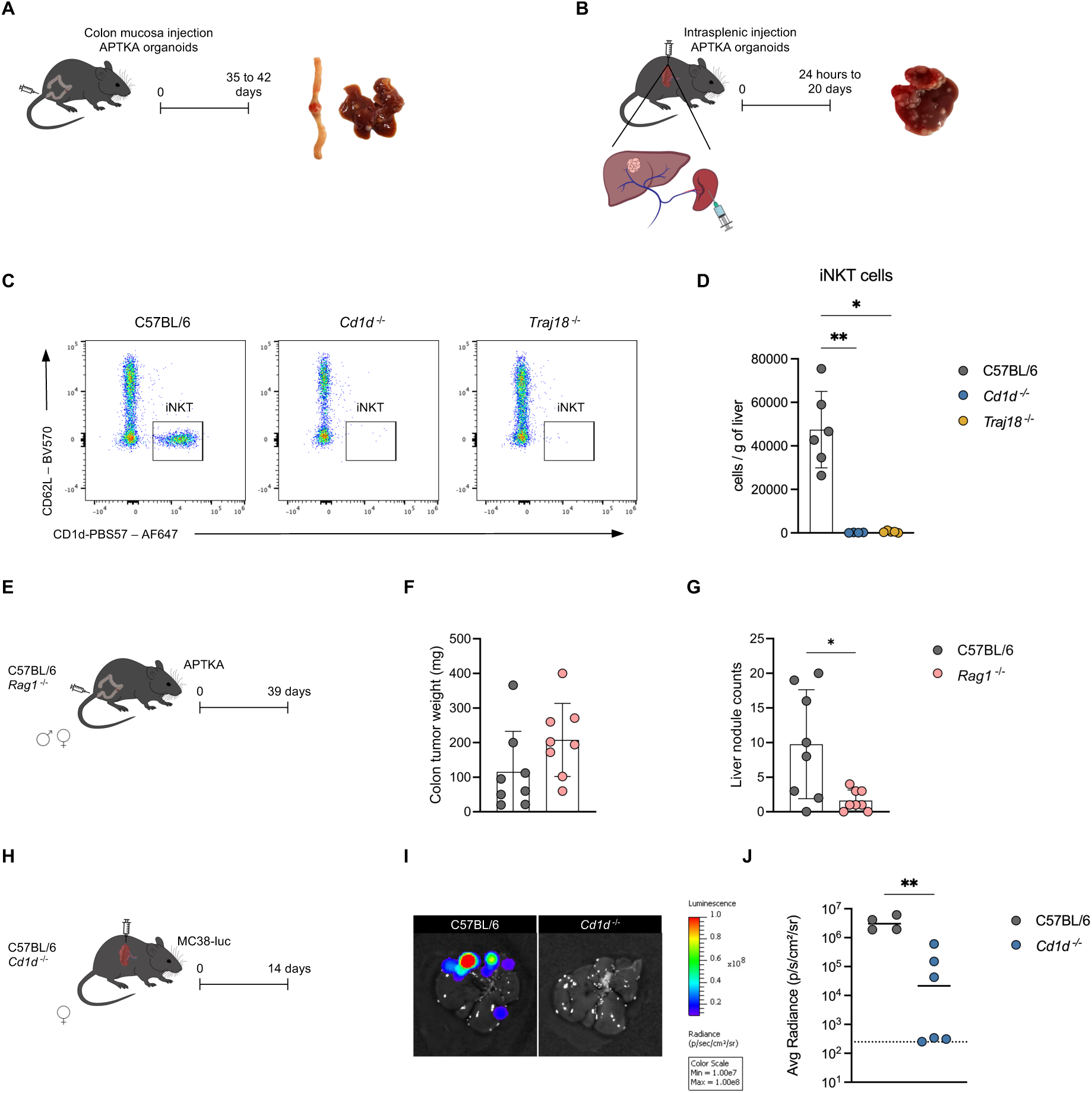
iNKT cells promote colorectal cancer-derived liver metastasis. (A) Orthotopic model of colorectal liver metastasis. (B) Intrasplenic model of colorectal liver metastasis. (C) Representative dot plots and (D) quantification of liver iNKT (CD1d-PBS-57^+^, CD62L^-^) cells. Samples were pre-gated on single, live, CD45^+^, lin^-^, TCRβ^+^, CD11b^-^ cells. Lin = Ly6G, CD19, CD115 and F4/80. Bars represent means ± SD. Symbols represent values from individual mice. \**p* < 0.05, \*\**p* < 0.01 (one-way ANOVA). (E) APTKA organoids were orthotopically injected into male and female C57BL/6 and *Rag1* ^-/-^ mice. Primary colon tumors and livers were collected and analyzed 39 days later. (F) Weight of primary colon tumor. (G) Quantification of liver metastasis. Bars represent means ± SD. symbols represent values from individual mice. \**p* < 0.05 (unpaired Mann-Whitney U test). (H) MC38-luc organoids were intrasplenically injected into male and female C57BL/6 and *Cd1d* ^-/-^ mice. *Ex vivo* bioluminescence of the liver was measured 14 days later. (I) Representative measurement and (J) quantification of liver bioluminescence. The dotted line represents the limit of detection. The solid lines represent the median. Symbols represent values from individual mice. \*\**p* < 0.01 (one-way ANOVA or unpaired Mann-Whitney U test).

**fig. S2.**
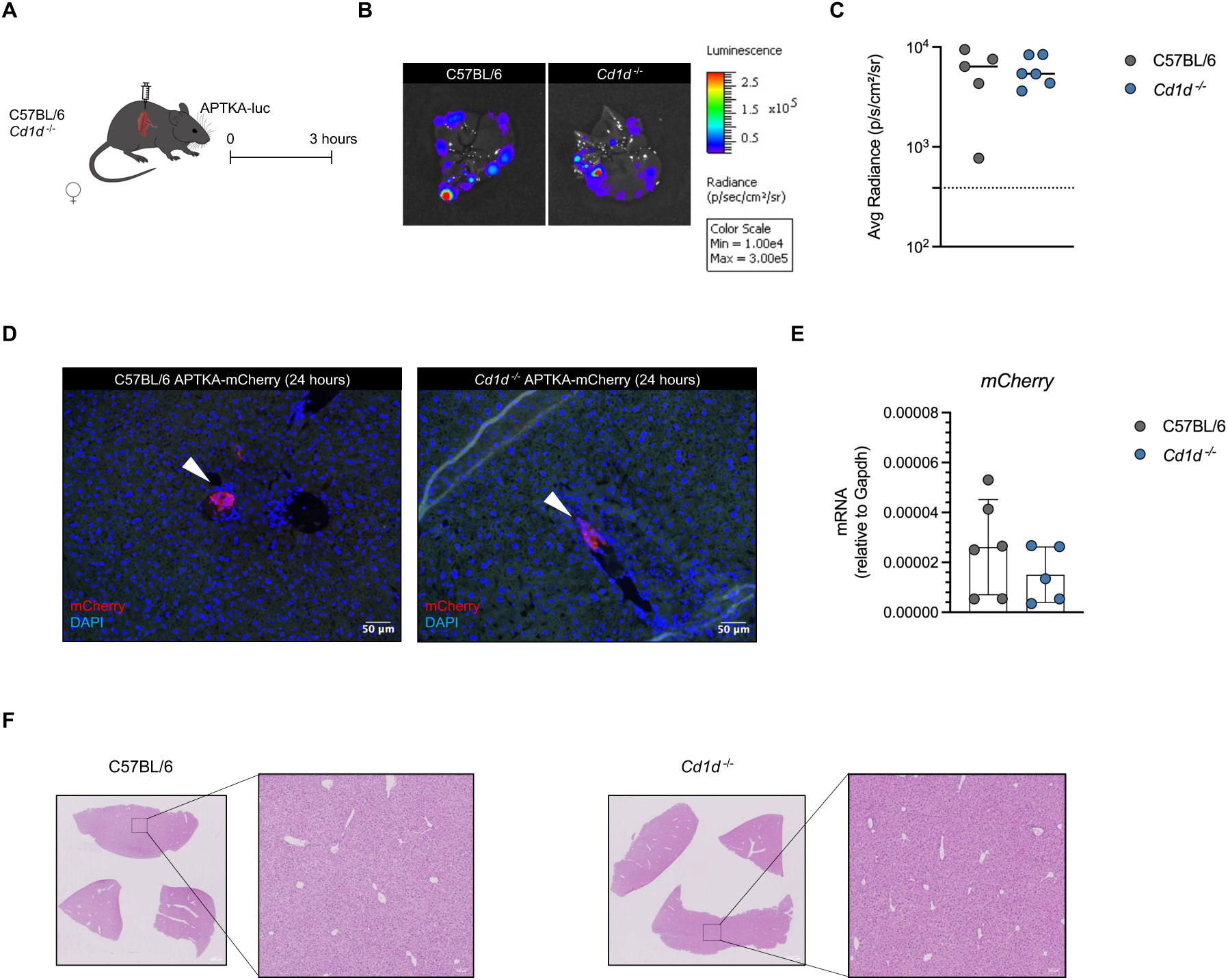
Lack of NKT cells does not affect the initial seeding of disseminating cancer cells to the liver. (A) APTKA-luc organoids were intrasplenically injected into female C57BL/6 and *Cd1d*^-/-^ mice. *Ex vivo* bioluminescence of the liver was measured 3 hours later. (B) Representative measurement and (C) quantification of liver bioluminescence. The dotted line represents the limit of detection. The solid lines represent the median. Symbols represent values from individual mice (unpaired Mann-Whitney U test). (D) Representative immunofluorescence image of disseminated cancer cells in C57BL/6 and *Cd1d* ^-/-^ mice. Arrows indicate APTKA-mCherry cells. (E) Relative quantification of mCherry transcripts in the liver lysate. Bars represent means ± SD, symbols represent values from individual mice (unpaired Mann-Whitney U test). (F) H&E-stained section of naïve livers from C57BL/6 and *Cd1d* ^-/-^ mice.

**fig. S3.**
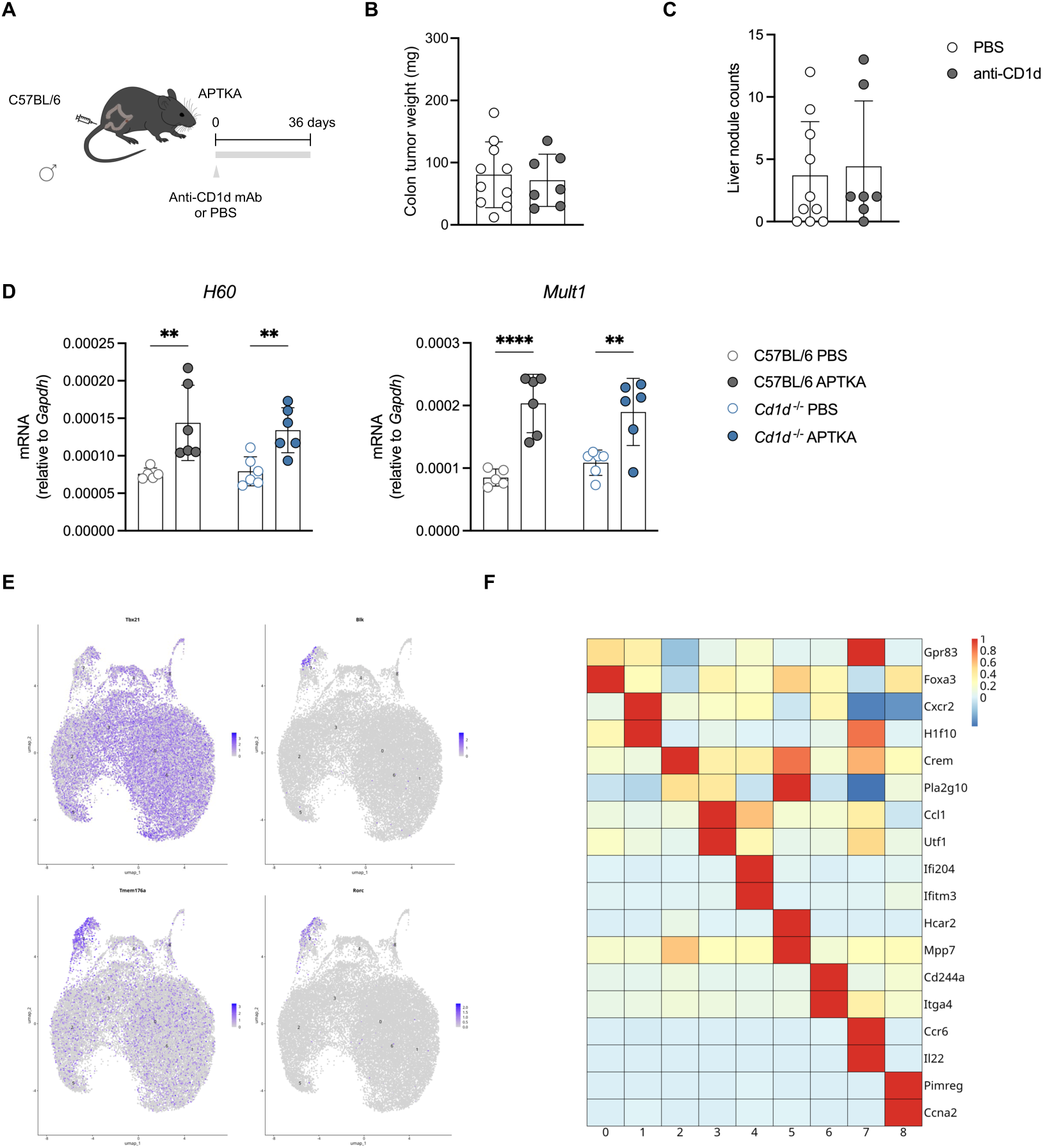
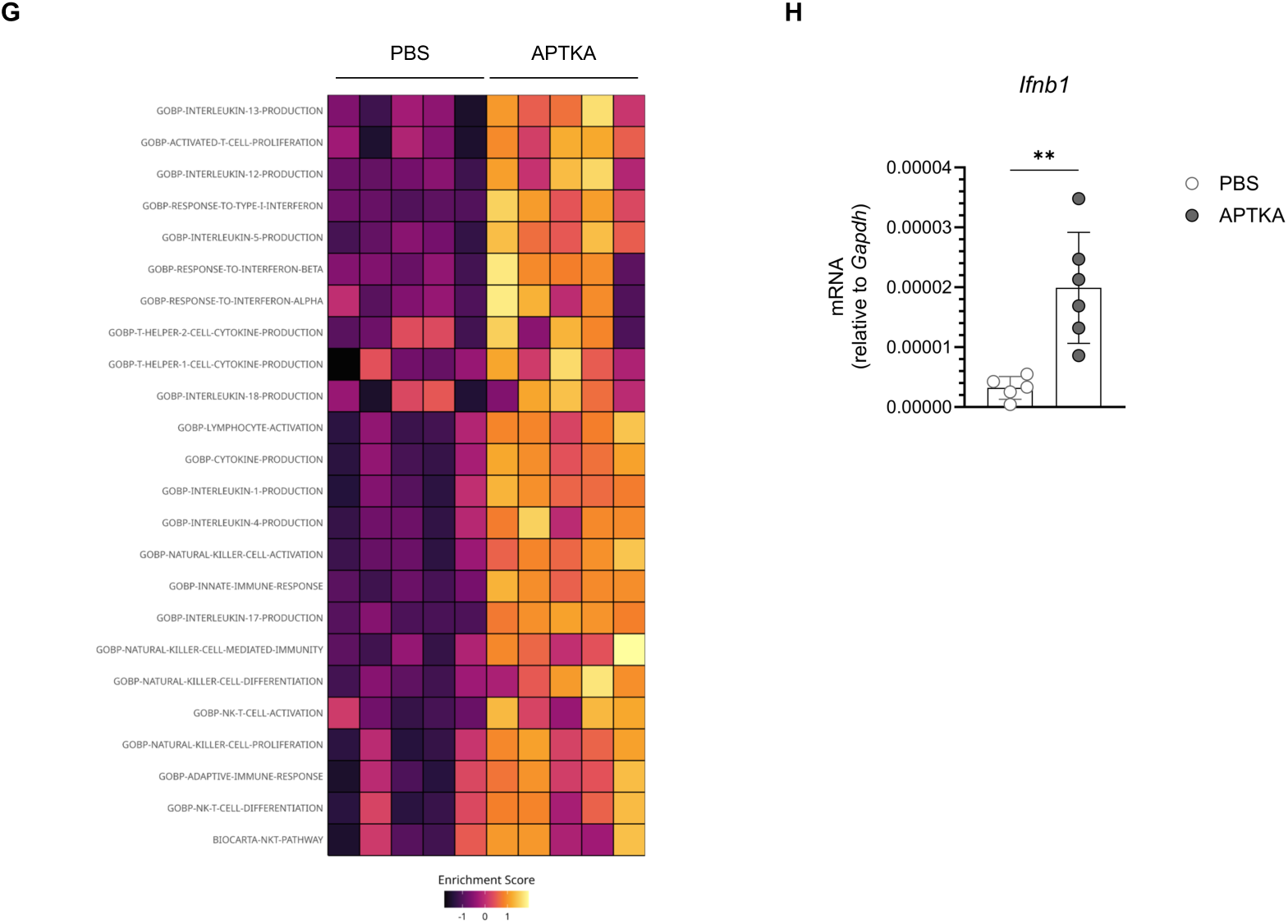
Cancer cell dissemination induces activation of hepatic iNKT cells independently of TCR signaling. (A) APTKA organoids were orthotopically injected into male C57BL/6 mice. Immediately thereafter, mice were injected i.p. with 100 μg of anti-CD1d blocking antibody or PBS. Anti-CD1d and PBS were administered twice per week for the duration of the experiment. Primary colon tumors and livers were collected and analyzed 36 days later. (B) Weight of primary colon tumors. (C) Quantification of liver metastasis. Bars represent means ± SD, symbols represent values from individual mice (unpaired Mann-Whitney U test). (D) Relative quantification of *H60* and *Mult1* transcripts in liver lysate 24 hours after intrasplenic injection of PBS or APTKA-mCherry in C57BL/6 and *Cd1d* ^-/-^ mice. Bars represent means ± SD, symbols represent values from individual mice \*\**p* < 0.01, *p*\*\*\*\* < 0.0001 (two-way ANOVA). (E) UMAP projection of NKT cells overlaid by expression of selected genes. (F) Heatmap showing the two genes with the highest expression in each identified Seurat cluster. (G) z-transformed enrichment score of selected gene sets across samples. (H) Relative quantification of *Ifnb1* transcript in liver lysate 24 hours after intrasplenic injection of PBS or APTKA-mCherry in C57BL/6 mice. Bars represent means ± SD, symbols represent values from individual mice \*\**p* < 0.01 (unpaired Mann-Whitney U test).

**fig. S4.**
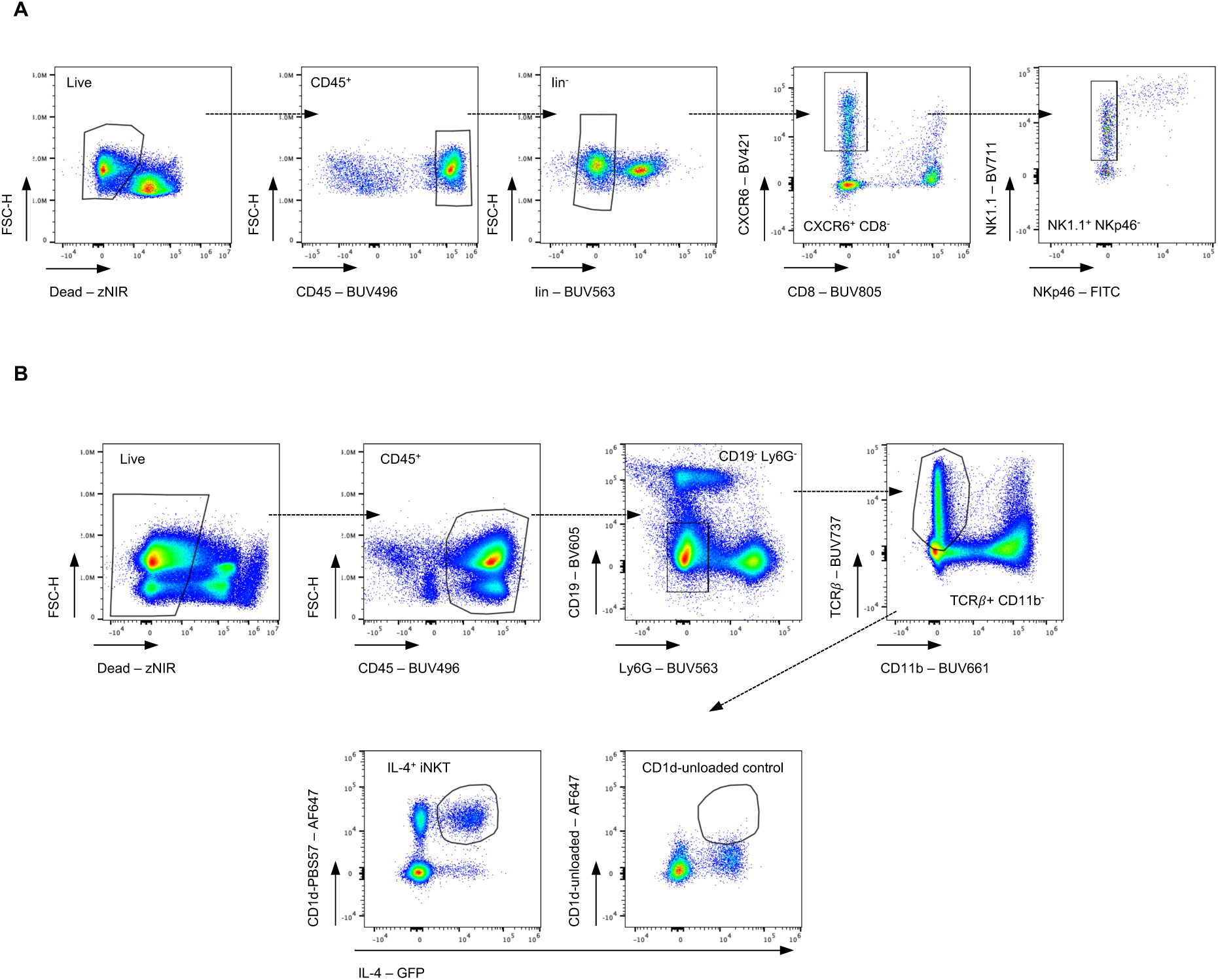
NKT cell gating strategy for sorting and flow cytometric analysis. (A) Gating strategy for sorting hepatic NKT cells. Samples were pre-gated on single cells. (B) Gating strategy used to identify hepatic iNKT cells in PBS or APTKA-injected *Il4 ^4G^*^et/wt^ mice. Samples were pre-gated on single cells.

**Table S1.**
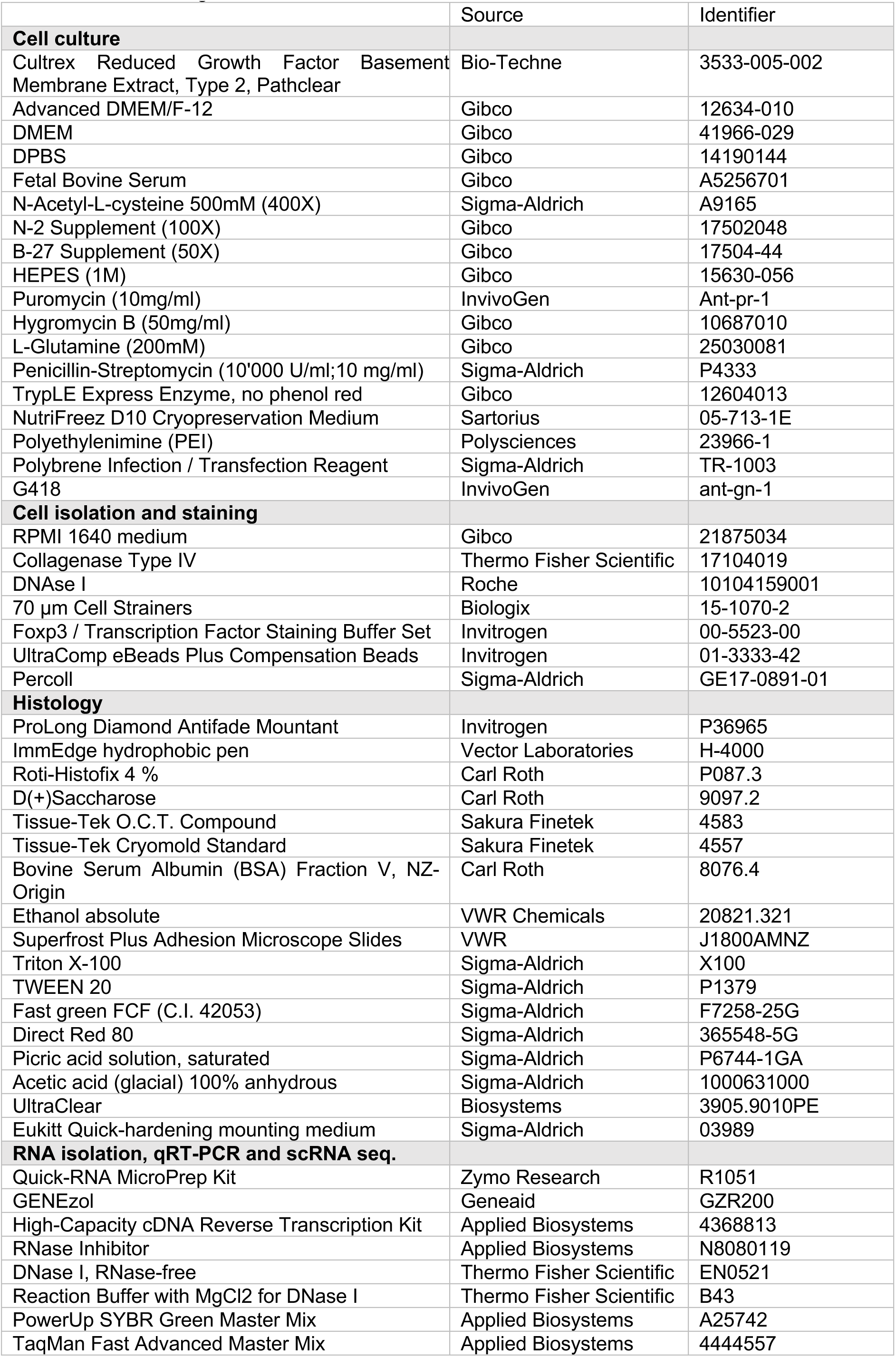
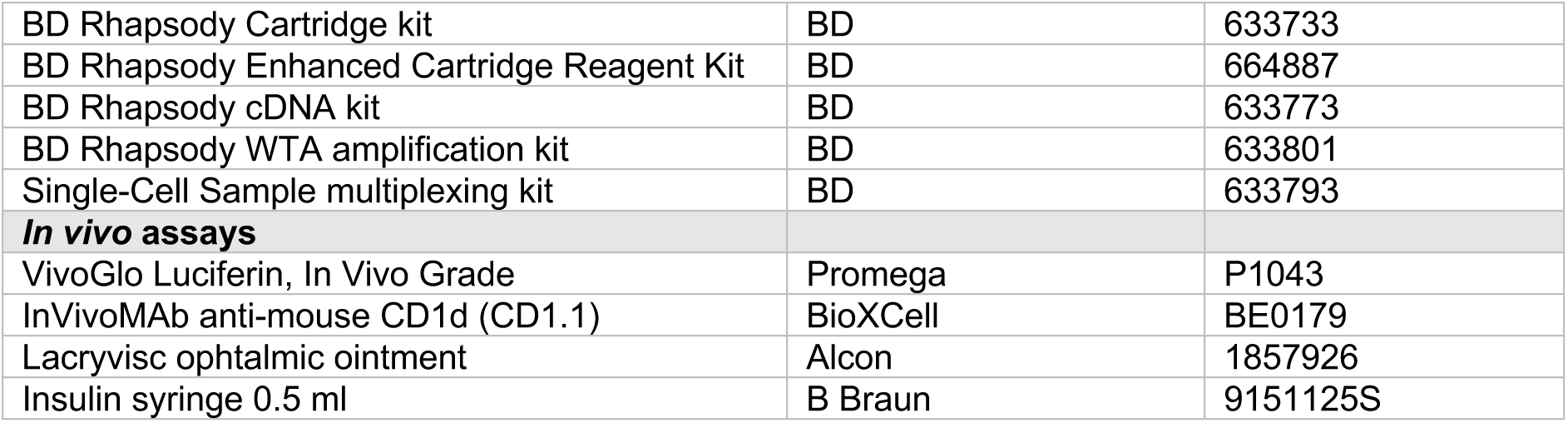
Kits and reagents.

**Table S2.** Special equipment.

|  |  |  |
| --- | --- | --- |
| **Colonoscopy-mediated injection** |  |  |
| Injection syringe | Hamilton | 7656-01 |
| Hopkins Telescope | Karl Storz | 64301AA |
| Intravascular catheter | BD | 381337 |
| Mucosal injection needle (33 gauge, small-hub RN needle, 45°, 400mm, point style 4) | Hamilton | 7803-05 |
| RN Nut | Hamilton | 30902 |
| Camera control unit IMAGE1 S CONNECT | Karl Storz | TC200 |
| Monitor | Karl Storz | 9826NB |
| Camera Head | Karl Storz | TH102 |
| Light cable | Karl Storz | 495 FS |
| Light source | Karl Storz | 20133720-1 |
| Operating sheath | Karl Storz | 61029D |
| 20-ml syringe | B Braun | 4606205V |
| **Intrasplenic injection** |  |  |
| Coated VICRYL Polyglactin 910 | Ethicon | V303H |
| Wound clips 9 mm | Clay Adams | 427631 |
| Wound clip applier | Fine Science tools | 12020-09 |
| **Miscellaneous** |  |  |
| BD FACS Aria III cell sorter | BD |  |
| Cytek Aurora 5L | Cytek Biosciences | N/A |
| LightCycler 480 II | Roche | 05015278001 |
| FastPrep-24 homogenizer | MP Biomedicals | 116004500 |
| Zirconium oxide beads | Labgene Scientific | BER103BK |
| Vectra 3 Automated Quantitative Pathology Imaging System | PerkinElmer | N/A |
| IVIS Imaging System 200 | PerkinElmer | N/A |
| Cryostar NX70 Cryostat | Epredia | 15380755 |

**Table S3.**
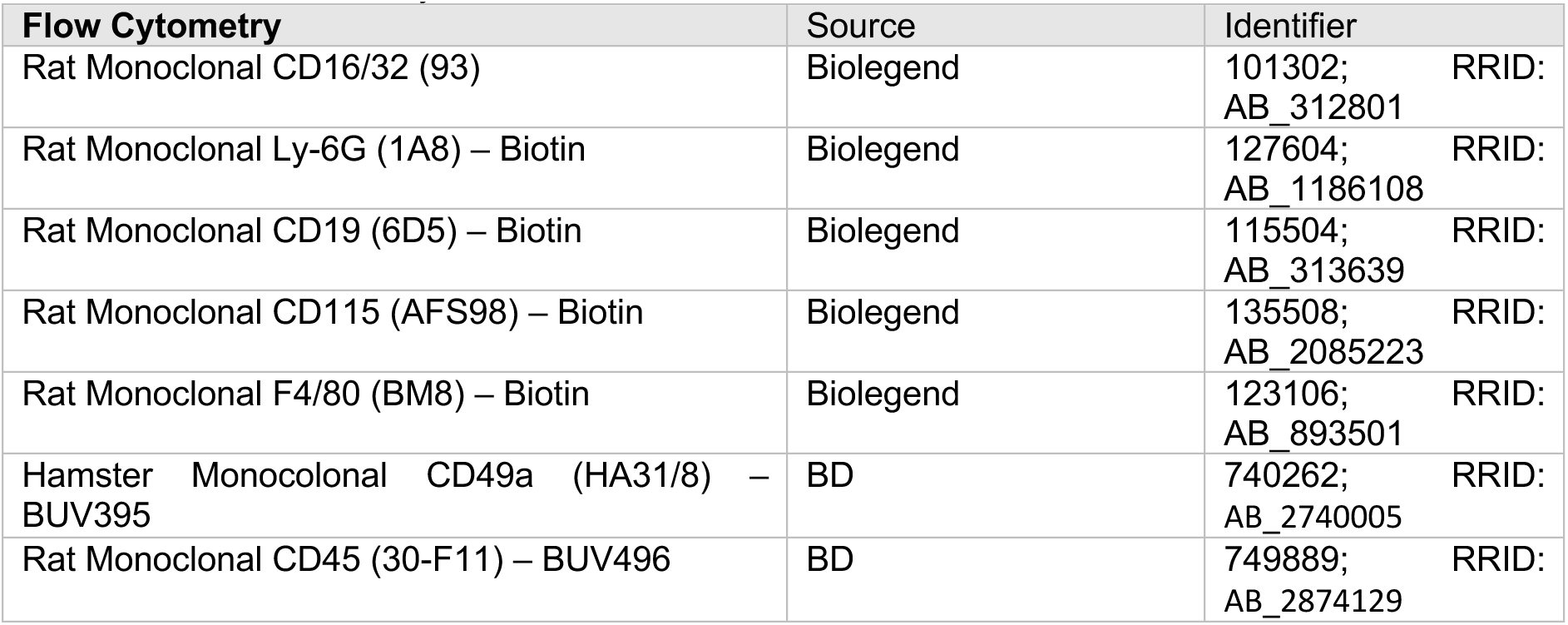
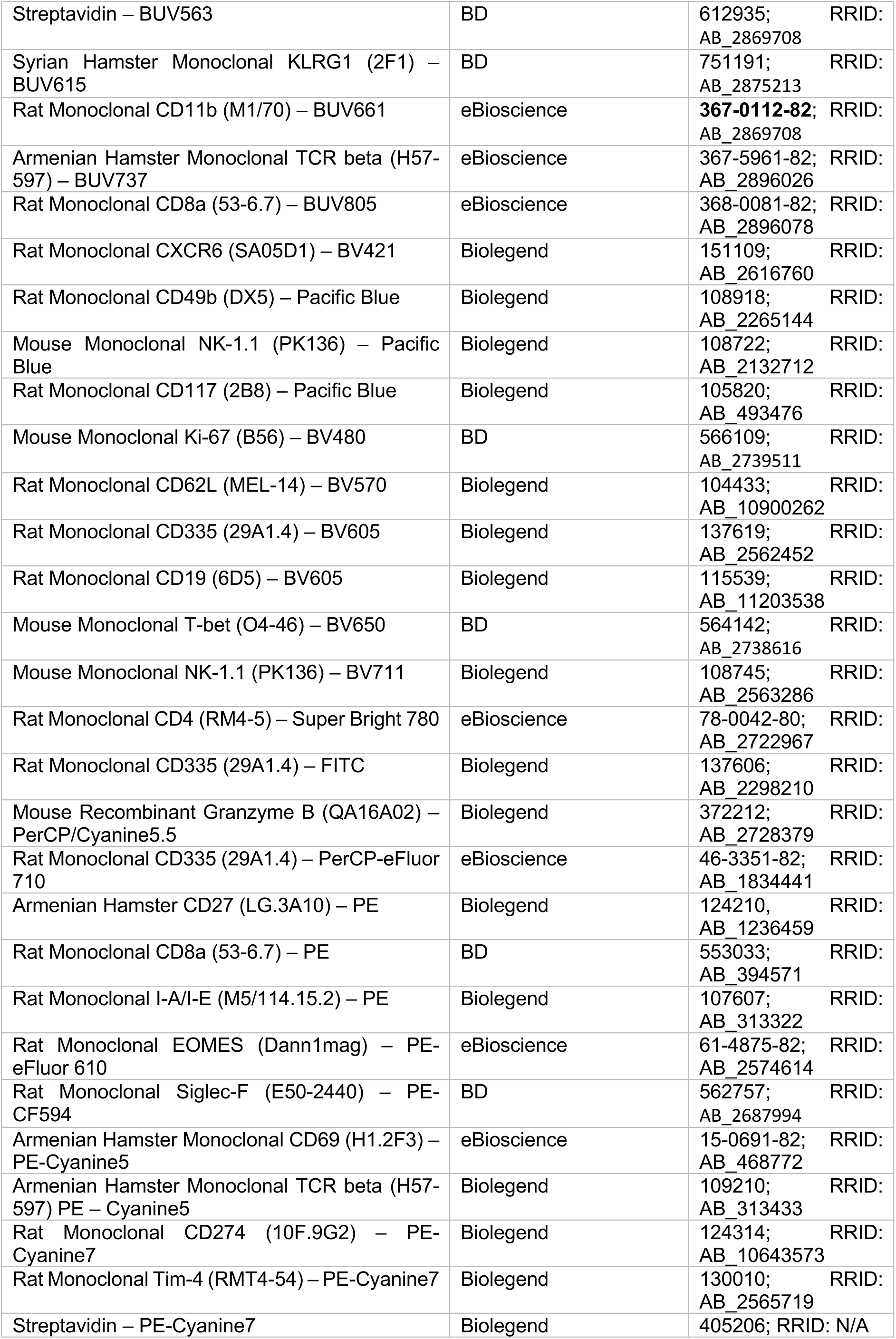
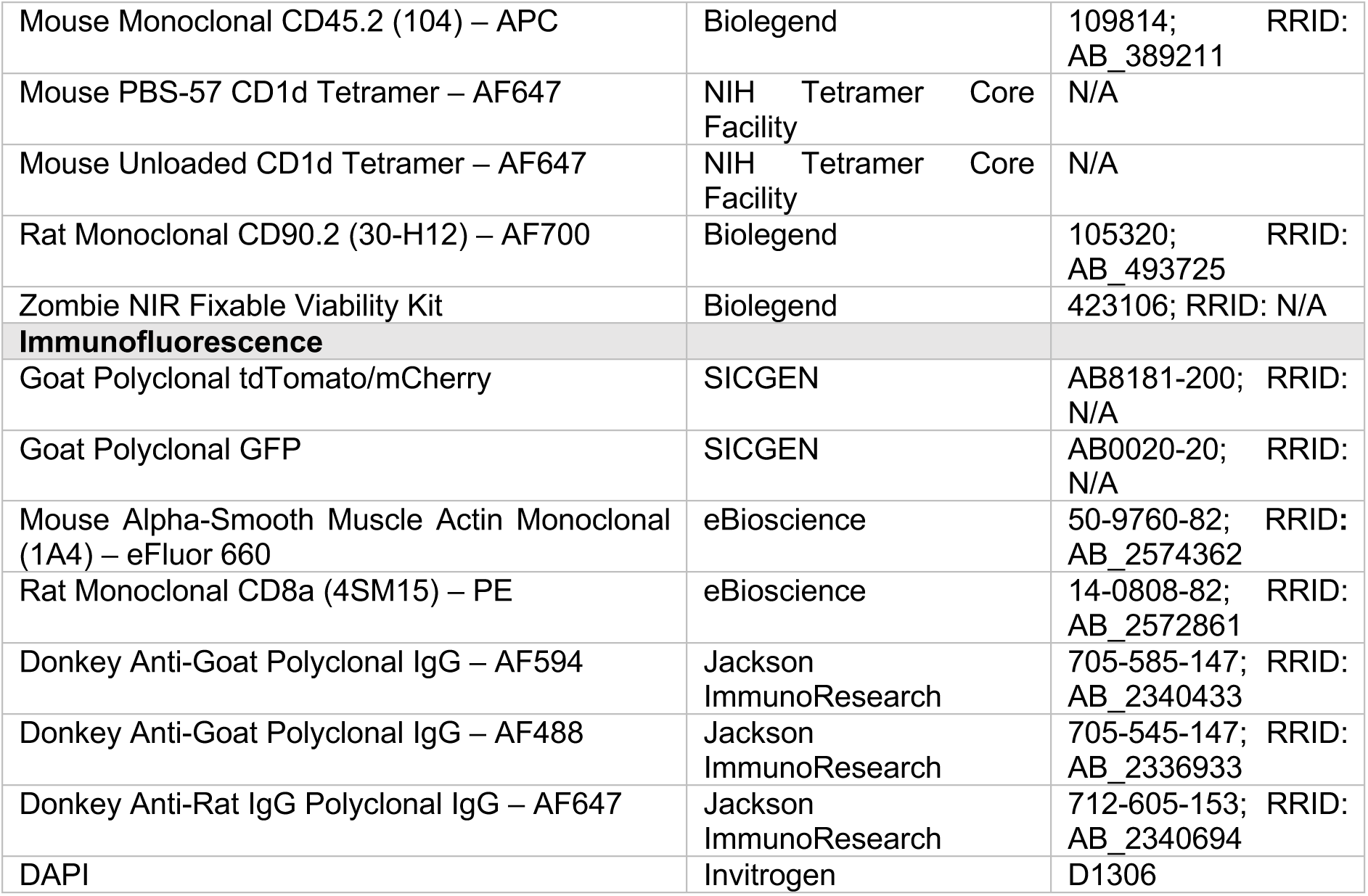
Antibodies and dyes.

**Table S4.**
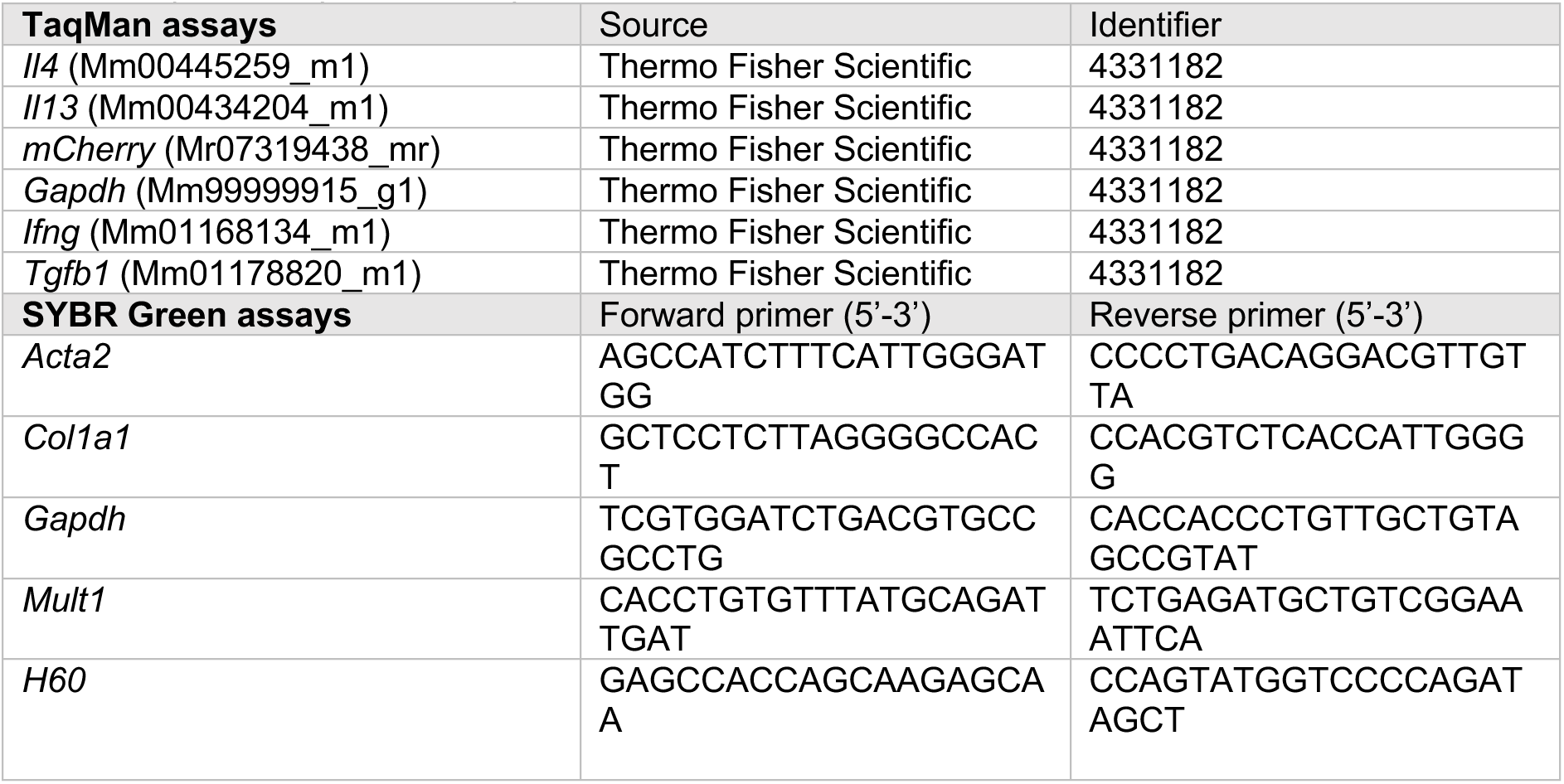
qRT-PCR primers and probes.

